# Genetic Landscape of Electron Transport Chain Complex I Dependency in Acute Myeloid Leukemia

**DOI:** 10.1101/513887

**Authors:** Irène Baccelli, Yves Gareau, Bernhard Lehnertz, Stéphane Gingras, Jean-François Spinella, Alexandre Beautrait, Sophie Corneau, Nadine Mayotte, Isabel Boivin, Simon Girard, Tara MacRae, Mélanie Frechette, Koryne Leveillé, Jana Krosl, Clarisse Thiollier, Vincent-Philippe Lavallée, Evgeny Kanshin, Thierry Bertomeu, Jasmin Coulombe-Huntington, Corinne St-Denis, Marie-Eve Bordeleau, Geneviève Boucher, Philippe P. Roux, Sébastien Lemieux, Mike Tyers, Pierre Thibault, Josée Hébert, Anne Marinier, Guy Sauvageau

## Abstract

Inhibition of oxidative phosphorylation (OXPHOS) is a promising therapeutic strategy in Acute Myeloid Leukemia (AML), but patients respond heterogeneously. Through chemically interrogation of 200 sequenced specimens, we identified Mubritinib as a strong *in vitro* and *in vivo* anti-leukemic compound, acting through ubiquinone-dependent inhibition of Electron Transport Chain complex I (ETC1). ETC1 targeting showed selective toxicity against a subgroup of chemotherapy-resistant leukemias exhibiting OXPHOS hyperactivity, high expression of mitochondrial activity-related genes, and mutations affecting *NPM1, FLT3* and *DNMT3A*. Altogether, our work thus identifies a novel ETC1 inhibitor with high clinical potential and reveals the landscape of OXPHOS dependency in AML.

---

Acute Myeloid Leukemia (AML) is a highly lethal disease, with a five-year overall survival rate of only 27% ^1,2^. Standard treatment for AML includes a combination of cytarabine (AraC) and anthracycline as an induction regimen, followed by consolidation chemotherapy or hematopoietic stem cell transplantation, depending on the patient’s genetic risk class ^3^. Although 60% to 70% of patients enter complete remission after induction regimen, most of them relapse within 3 years, due to the outgrowth of therapy resistant AML leukemic stem cells (LSCs) ^4,5^. Thus, the identification of novel treatment strategies, in particular for poor outcome AML patients, represents an outstanding medical need.

The development of novel therapeutic approaches in AML has long been precluded by the absence of culture conditions that preserve the activity of LSCs *in vitro*. Recently, our group developed a culture method which maintains LSC activity for several days ^6^, thus enabling relevant cell-based chemical interrogation of the disease ^7–12^. Nevertheless, normal and leukemic stem cells share numerous biological traits, making their specific eradication challenging ^13^. For instance, the initially described CD34^+^/CD38^-^ LSC cell surface phenotype ^5^ also characterizes normal hematopoietic stem cells (HSCs). Furthermore, although LSC-specific gene expression signatures are being developed ^14^, their transcriptional landscape appears to be highly reminiscent of that of HSCs ^15^, probably because LSCs are oftentimes derived from HSCs ^4,16^.

However, striking differences in energy metabolism between normal and leukemic stem cells have recently been revealed. HSCs rely primarily on anaerobic glycolysis rather than mitochondrial oxidative phosphorylation (OXPHOS) for energy production and repression of mitochondrial metabolism by autophagy is in fact required for their long-term self-renewing capacity ^17–19^. In stark contrast, AML LSC protein expression profiles are enriched for hallmarks of OXPHOS ^20^ and LSCs rely on mitochondrial function for their survival: they are sensitive to tigecycline, an antibiotic that inhibits mitochondrial protein synthesis ^21^ and to 2’3’-dideoxycytidine (ddC), a selective inhibitor of mitochondrial DNA replication ^22^. Furthermore, AML cells appear to be overall characterized by high OXPHOS activity and high mitochondrial mass, accompanied by low respiratory chain spare reserve capacity as compared to their normal hematopoietic counterparts ^23^, and are sensitive to inhibition of the mitochondrial protease ClpP ^24^. Interestingly, AML cells resisting cytarabine treatment in mouse xenografts were recently reported as relying primarily on OXPHOS for their survival ^25^ and a novel NADH dehydrogenase inhibitor shows promising anti-leukemic activity ^26^. Last but not least, OXPHOS suppression induced by the combination of BCL-2 inhibitor venetoclax and azacytidine selectively targets LSCs and results in deep and durable remissions in AML patients ^27,28^. Hence, inhibition of mitochondrial function represents a promising therapeutic strategy in AML. While a large portion of AMLs exhibit strong sensitivity to inhibition of mitochondrial metabolism, resistance is frequently observed ^29,30^. However, the determinants of resistance and sensitivity to OXPHOS targeting in this genetically highly heterogeneous disease remain so far unclear. In this study, we interrogated 200 genetically diverse primary AML patient specimens of the well-characterized Leucegene cohort (www.leucegene.ca) using an unbiased chemo-genomic approach. Through this process, we identified a novel ubiquinone-dependent Electron Transport Chain (ETC) complex I inhibitor and uncovered the genetic landscape of OXPHOS dependency in this disease.

## Results

### Mubritinib targets a subset of poor outcome AMLs

The Leucegene collection of sequenced AML specimens comprises 263 *de novo* non-M3 samples, originating from patients who received intensive standard chemotherapy treatment, and for whom long-term survival data is available (termed “prognostic cohort”). Within this cohort, we first compared the clinical and genetic characteristics of specimens from “poor outcome” patients (hereafter defined by an overall-survival strictly below 3 years, n=183) to that of specimens from “good outcome” patients (survival≥3 years, n=80).

As expected, the poor outcome group comprised most (87%) adverse cytogenetic risk specimens and a minority (34%) of favorable risk AMLs, but it also included the vast majority (73%) of intermediate cytogenetic risk samples, highlighting the heterogeneity of survival rates within this latter class (Fig. S1a). Accordingly, the poor outcome cohort was overall strongly enriched for adverse cytogenetic risk AMLs (p=0.0009), whereas the good outcome cohort associated with favorable cytogenetic risk specimens (p<0.0001), including Core Binding Factor (CBF) leukemias (p=4x10^-8^, Fig. S1b). In agreement with other reports ^31^, we noted an increase in patient age (p<0.0001), relapse rates (p=5x10^-12^, Fig. S1b), and frequencies of lesions affecting *TP53* (p=0.0075, Fig. S1c) in the poor outcome group. Also in line with other studies ^32–34^, poor outcome AMLs strongly associated with overexpression of Homeodomain-containing transcription factor *HOXA9* and other *HOX-*network genes. Interestingly, genes overexpressed more than 10-fold in poor outcome AMLs all belonged to the *HOXA* cluster (Fig. S1d).

Next, in order to uncover new therapeutic targets for poor outcome AML patients, we chemically interrogated primary specimens of the Leucegene prognostic cohort originating from poor and good outcome patients. Using LSC-activity maintaining culture conditions ^6^ and a panel of inhibitors targeting receptor tyrosine kinases, as well as components of the RAS and PI3K pathways, we first compared the Growth Inhibitory 50 (GI50) values of these compounds in a small cohort of primary AMLs (Fig. 1a, patient characteristics in Table S1). Mubritinib, a compound described as a specific ERBB2 inhibitor ^35^, appeared as the most selective inhibitor towards specimens belonging to the poor outcome group (p=0.009, Fig. 1b). In order to validate this observation, we then assessed the sensitivity to Mubritinib treatment in a large and heterogeneous cohort (Fig. 1c, n=200, patient characteristics in Table S2). This secondary screen confirmed that leukemic cells originating from poor outcome patients are more sensitive to Mubritinib than those from good outcome patients (p=0.0048, Fig. 1d). Importantly, the proliferation of CD34^+^ cord blood control cells was not affected by Mubritinib treatment, for concentrations up to 10μM (n=5 unrelated samples, Fig. 1d).

**Fig. 1.**
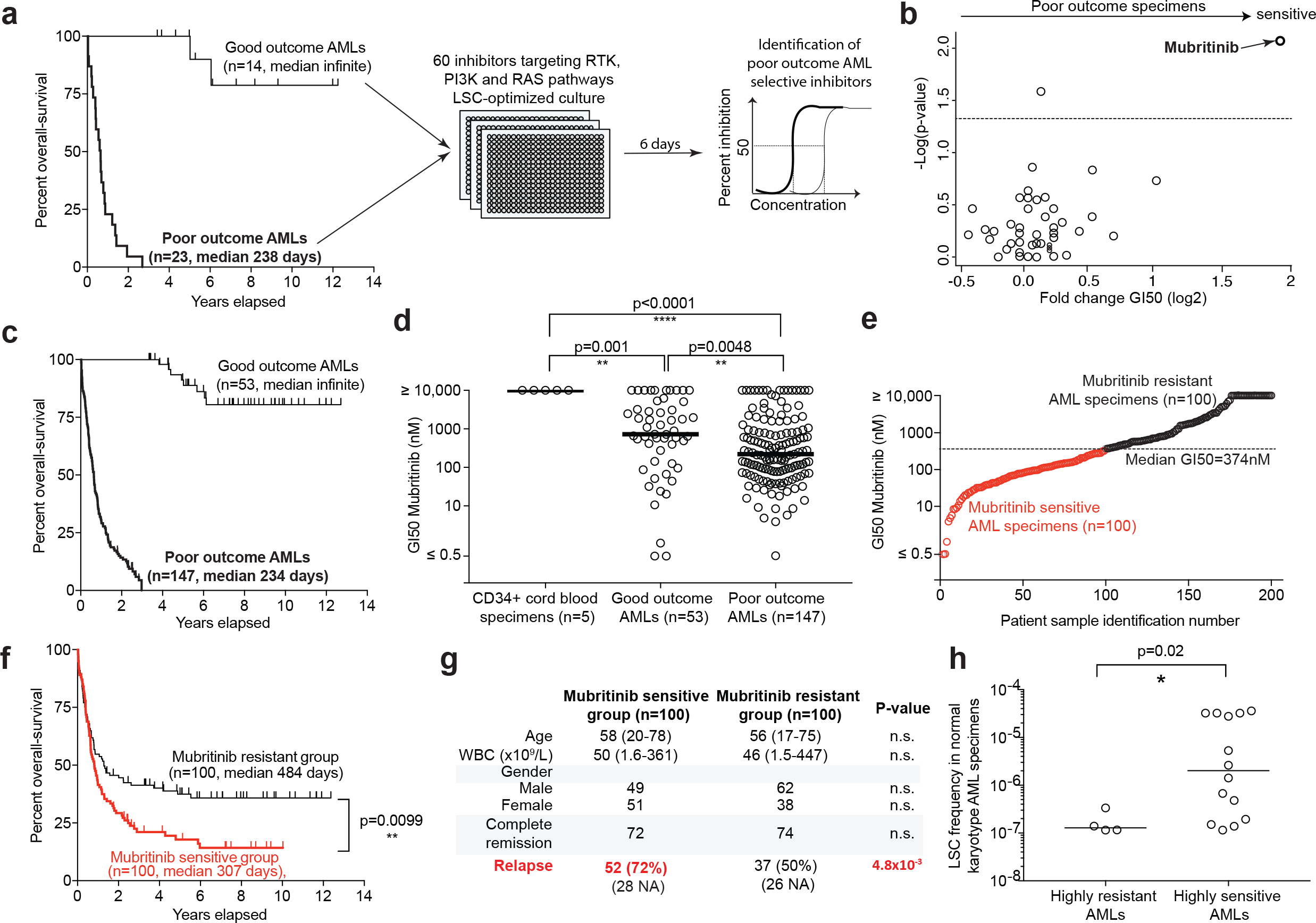
Identification of Mubritinib as a potent and selective inhibitor for poor outcome AML. **a**, Workflow and **b**, result of the primary screen (see also Table S1). **c**, Set-up and **d**, result of the validation screen (see also Table S2). **e**, Definition of Mubritinib sensitive and resistant groups. **f**, Overall surviva and **g**, general clinical features of patients belonging to the sensitive and resistant groups. **h**, LSC frequencies in highly sensitive (GI50 values within the lower tertile of the cohort) and highly resistant (GI50 values within the higher tertile of the cohort) normal karyotype AMLs. In **b**, the horizontal grey line corresponds to p=0.05. Statistical assessments were performed using the Mann-Whitney test (**b**, **d** and **h**), the log-rank test (**f**) and the Fisher’s exact test (**g**). Data in **d** and **h** are represented as median values.

Mubritinib GI50 values varied widely among AML specimens (median: 374nM, Fig. 1e). We first compared the general features of Mubritinib sensitive (GI50 below median, n=100) and resistant specimens (GI50 above median, n=100, Fig. 1e). Importantly, we detected no significant difference between the *in vitro* proliferation rates of untreated specimens from both groups (Fig. S2a), indicating that the differences in GI50 values were not due to a proliferation bias. Patients belonging to the Mubritinib sensitive group exhibited decreased overall-survival rates compared to patients belonging to the resistant category (p=0.0099, Fig. 1f). While we observed no difference in complete remission rates between patients belonging to the Mubritinib sensitive and resistant classes, sensitivity associated with a significant increase in relapse rates (p=4.8x10^-3^, Fig. 1G). In line with these results, within the relatively homogenous normal karyotype AML genetic subtype, leukemias highly sensitive to Mubritinib (GI50 values in the lower tertile) exhibited elevated LSC frequencies as compared to highly resistant specimens (GI50 values in the upper tertile, p=0.02, Fig. 1h).

We next investigated the clinical and mutational characteristics of Mubritinib resistant and sensitive AMLs (Fig. 2a and Table S3). Overall, resistance to Mubritinib associated with the favorable cytogenetic risk group (p=4.4x10^-8^), with the inv(16) subtype (p=2.9x10^-5^) and more generally with CBF AMLs (p=4x10^-8^). In addition, resistance to Mubritinib associated with the presence of *KIT* mutations (p=4.2x10^-4^) and more generally with mutations affecting the RAS-MAPK signaling pathway (p=1.8x10^-5^, Fig. 2a-c and Table S3). Conversely, sensitivity to Mubritinib strongly associated with the intermediate cytogenetic risk category (p=5.6x10^-8^), with normal karyotype specimens (p=8.9x10^-7^), with the presence of mutations affecting *NPM1* (p=3.5x10^-5^), *FLT3-*ITD (p=7.2x10^-5^), and *DNMT3A* (p=4.1x10^-7^), as well as with mutations involving genes belonging to the DNA methylation class (comprising *DNMT3A, IDH1, IDH2* and *TET2*, p=4.1x10^-8^, Fig. 2a-c and Table S3). Interestingly, the recently identified adverse outcome triple mutated AMLs ^36,37^ (carrying *NPM1*, *FLT3*-ITD and *DNMT3A* co-occurring mutations, n=34 out of 200) were significantly enriched within the sensitive group (p=0.002, Fig. 2d) and exhibited high sensitivity to Mubritinib compared to other samples (median GI50=96nM *versus* 490nM for all other AMLs, p=0.0002, Fig. S2b).

**Fig. 2.**
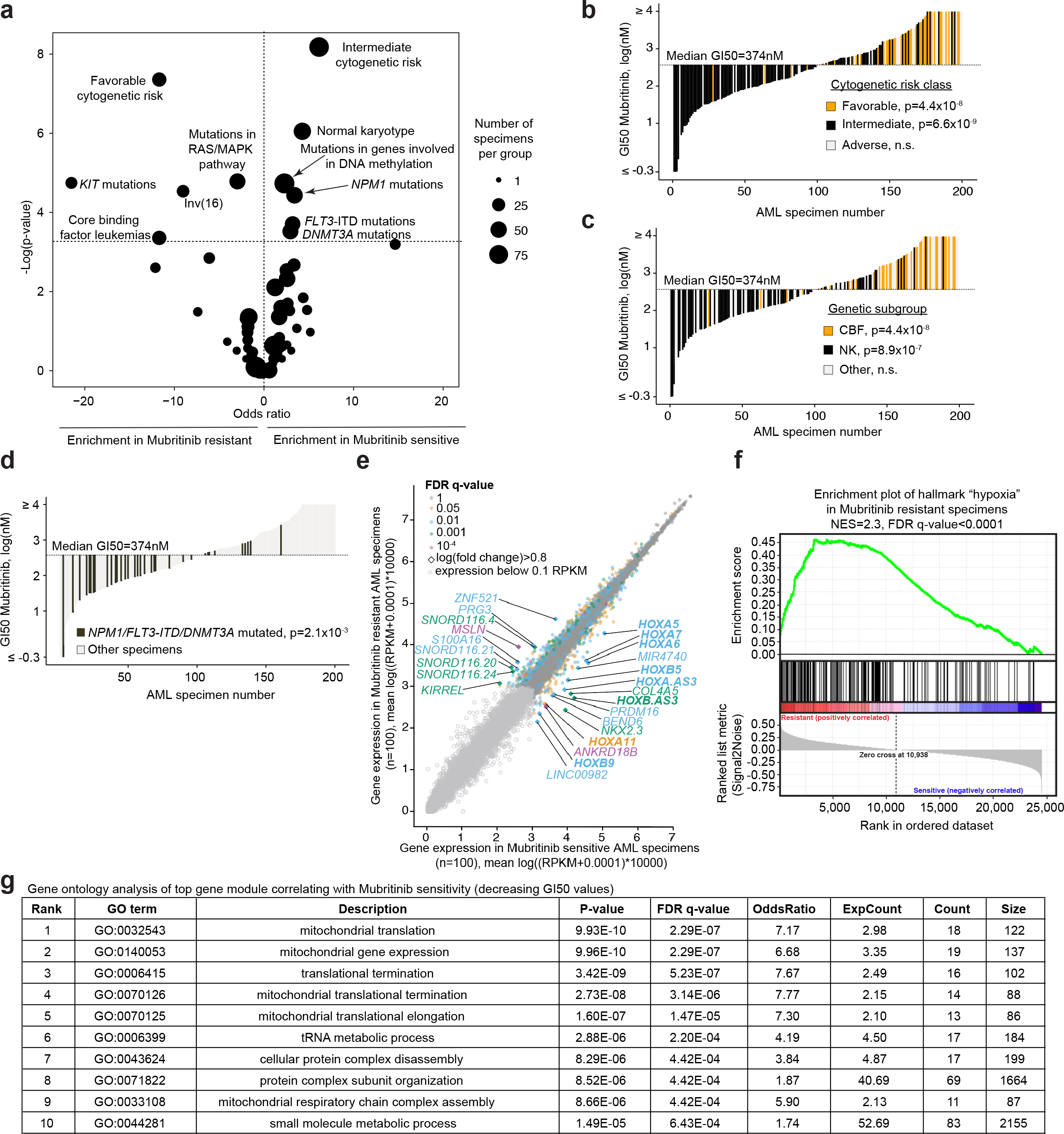
Clinical, mutational and transcriptional landscape of Mubritinib sensitivity in AML. **a**, Clinical and mutational features enriched in Mubritinib sensitive and resistant AML groups (see definition of groups in Fig. 1e and corresponding data in Table S3). Mubritinib GI50 values according to **b**, cytogenetic risk classes, **c**, genetic subgroups and **d**, the presence of co-occurring *FLT3*-ITD, *DNMT3A* and *NPM1*-mutations (see also Figure S2b). **e**, Most differentially expressed genes between Mubritinib-resistant and -sensitive specimens, highlighting *HOX*-network genes in bold (see also Table S4). **f**, Result of gene set enrichment analysis comparing most sensitive (lower quartile, n=50) to most resistant (upper quartile, n=50) specimens. G. Gene ontology analysis of top gene module anti-correlating with Mubritinib GI50 values resulting from Weighted Gene Co-expression Network Analysis (WGCNA, see methods). Statistical assessments were performed using the Bonferroni-corrected Fisher’s exact test (**a-d**), and analysis of differential gene expression was performed using the Wilcoxon rank-sum test and the false discovery rate (FDR) method (**e**). For gene set enrichment analysis in **f**, 1,000 permutations were performed by gene set.

In addition, and similar to poor outcome specimens (Fig. S1d), comparison of transcriptomic signatures of Mubritinib sensitive and resistant leukemic samples highlighted *HOX-*network gene overexpression in sensitive specimens (bold characters in Fig. 2e and Table S4). Gene set enrichment analysis of most sensitive and resistant specimens (lower quartile *versus* upper quartile) revealed that resistant specimens display strong hallmarks of hypoxia (NES=2.32, FDR q-value<0.0001, Fig. 2f). In contrast, by weighted gene co-expression network analysis (WGCNA) ^38^, the module whose expression most significantly associated with Mubritinib sensitivity in the 200 tested specimens (p=0.03, ranked co-first with another module) was highly enriched for genes involved in mitochondrial activity. Indeed, gene ontology analysis revealed that five of the top ten enriched biological processes in this module relate to mitochondrial function, including mitochondrial respiratory chain complex assembly (GO:0033108, FDR q-value=0.0004, Fig. 2g).

### Mubritinib impairs mitochondrial respiration

Mubritinib has been reported as a specific ERBB2 inhibitor ^35^. Unexpectedly, none of the AML patient specimens tested in the primary screen (Fig. 1a and Table S1) responded to Lapatinib, another potent ERBB2 inhibitor ^39^ (Fig. 3a). In contrast to healthy and cancer breast tissues, *ERBB2* mRNA expression levels were very low in both Mubritinib sensitive and resistant AMLs (average 0.6 RPKM in both groups, Fig. S3a). Moreover, ERBB2 protein expression could not be detected by flow cytometry in the Mubritinib sensitive *DNMT3A-* and *NPM1-*mutated human AML cell line OCI-AML3 ^40,41^ (Fig. 3b and Fig. S3b) or in Mubritinib-sensitive AML specimens (Fig. 3b). The same observation was made by liquid chromatography-mass spectrometry (LC-MS/MS) analysis of OCI-AML3 cells (Table S5). Taken together, these data indicate that, in the context of AML, Mubritinib’s activity is likely not mediated by ERBB2 inhibition.

**Fig. 3.**
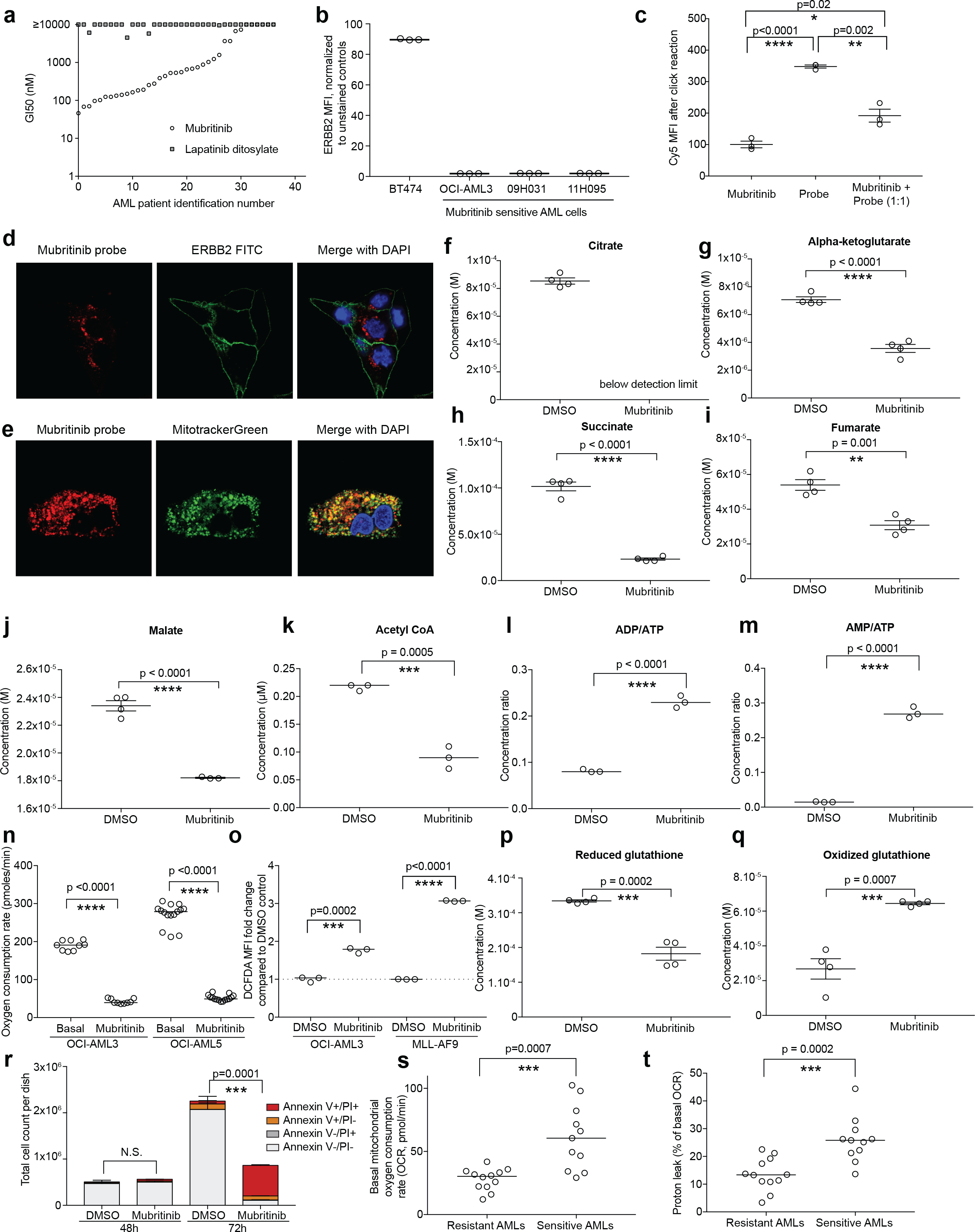
Mubritinib impairs mitochondrial respiration. **a,** Mubritinib and Lapatinib GI50 values in AML specimens (see also Table S1). **b**, ERBB2 protein expression levels (Mean Fluorescent Intensity, MFI). **c**, Validation of the specificity of the Mubritinib alkyne probe in OCI-AML3 cells (see also Fig. S3c). Confocal microscopy imaging of signals induced by **d**, the Mubritinib alkyne probe and an ERBB2 antibody as well as by **e**, the Mubritinib alkyne probe and the Mitotracker green dye in BT474 cells. Effect of Mubritinib treatment (500nM, 20h) on **f**, citrate, **g**, alpha-ketoglutarate, **h**, succinate, **i**, fumarate, **j**, malate, **k**, AcetylCoA, **l**, ADP/ATP ratio, **m**, AMP/ATP ratio concentrations in OCI-AML3 cells. Effect of acute (1μM) Mubritinib treatment on **n**, oxygen consumption rates (OCR) in 2 human AML cell lines. Effect of Mubritinib treatment (500nM, 20h) on **o**, 2’,7’-dichlorofluorescin diacetate (DCFDA) fluorescence intensity, as well as on **p**, reduced and **q**, oxidized glutathione concentrations in OCI-AML3 cells. **r**, Apoptotic cell death rates in OCI-AML3 cells upon Mubritinib treatment (500nM), (see also Fig. S3i). **s**, Basal mitochondrial OCR and **t**, proton leak rates measured in Mubritinib sensitive and resistant primary AML specimens (see also Fig. S3j-k and Table S6). Statistical assessments were performed using the unpaired two-tailed *t*-test. In **p**, the number of Annexin V and Propidium Iodide (PI) double positive cells were considered for statistical assessment. Data are represented as median values (**b** and **k-o**) or mean values of triplicates or quadruplicates, with SEM.

In order to understand Mubritinib’s mode of action, we first investigated which sub-cellular compartment it targets using a specific alkyne probe (Fig. 3c and Fig. S3c). We noted that, while the signal induced by the Mubritinib probe did not co-localize with the receptor tyrosine kinase ERBB2 (Fig. 3d), it co-localized with a mitochondrial dye (Fig. 3e), suggesting that Mubritinib accumulates in mitochondria. Furthermore, LC/MS analyses revealed consistent decrease of Krebs cycle metabolite concentrations (citrate, alpha-ketoglutarate, succinate, fumarate, malate and acetyl coA Fig. 3f-k) in treated OCI-AML3 cells, indicating that mitochondrial respiration is impaired. In line with these results, ADP/ATP and AMP/ATP ratios were largely increased upon treatment (Fig. 3l-m). In addition, monitoring of oxygen consumption rates in two different human AML cell lines with a Seahorse analyzer revealed that mitochondrial respiration is inhibited upon Mubritinib treatment (Fig. 3n). Concurrently, we measured increased extra-cellular acidification rates (ECAR) in Mubritinib treated cells, a phenomenon which could be rescued by 2-deoxy-D-glucose (2-DG, an inhibitor of glycolysis, Fig. S3d-e), suggesting an upregulation of glycolytic activity upon treatment. Accordingly, Mubritinib treated cells showed increased intracellular and extracellular lactate concentrations (Fig. S3f-g). Taken together, these data indicate that Mubritinib treatment of AML cells induces a switch from OXPHOS towards glycolytic metabolism.

In addition, 2’,7’ -dichlorofluorescin diacetate (DCFDA) staining of Mubritinib-sensitive human (OCI-AML3) or murine (MLL-AF9, Fig. S3h) AML cells revealed reactive oxygen species (ROS) accumulation upon Mubritinib treatment (Fig. 3o). LC/MS analyses highlighted decreased levels of reduced glutathione (Fig. 3p) concomitant with increased levels of oxidized glutathione (Fig. 3q) in presence of Mubritinib, confirming the induction of oxidative stress in treated cells. In response to treatment, sensitive AML cells underwent apoptotic death as assessed by flow cytometry using Annexin V and propidium iodide staining in OCI-AML3 cells (Fig. 3r) and MLL-AF9 cells (Fig. S3i). Last but not least, we monitored the OXPHOS activity profiles of genetically diverse primary AML specimens (see list in Table S6) that are either sensitive (n=11, GI50 < 100nM) or resistant (n=12, GI50> 5μM) to Mubritinib treatment. We found that basal mitochondrial oxygen consumption and proton leak rates are significantly higher in sensitive leukemias as compared to resistant AMLs, while extracellular acidification rates do not differ between the two groups of specimens (Fig. 3s-t and S3j-k). The results are in line with our observation that expression levels of genes involved in mitochondrial activity correlate with Mubritinib sensitivity Fig. 2g) and suggest that Mubritinib-sensitive AMLs are more heavily relying on OXPHOS for energy production than Mubritinib-resistant leukemias (Fig. 2g). In addition, as proton leak mediates a decrease in OXPHOS-induced ROS ^42^, these results also highlight a possible mechanism through which Mubritinib-sensitive AMLs are able to withstand such sustained basal OXPHOS hyperactivity.

### Mubritinib is a novel ETC complex I inhibitor

In order to further dissect Mubritinib’s mode of action, we next monitored changes in protein and phospho-protein abundance upon Mubritinib treatment using Stable Isotope Labeling by Amino acids in cell Culture (SILAC)-proteomic (Fig. 4a) and phospho-proteomic (Fig. 4b) approaches, respectively. In these experiments, pyruvate dehydrogenase (PDH) E3 subunit was the most prominent down-regulated protein in Mubritinib treated cells (Fig. 4a), while the inactivating phosphorylation ^43^ of serine 293 of PDH E1 subunit was markedly increased upon treatment (Fig. 4b). Although we could confirm the inhibitory effect of Mubritinib on PDH using a cell-based enzymatic activity assay (Fig. 4c, p=0.0098), we found that this effect was in fact indirect as Mubritinib (25μM, cell-free context) did not inhibit the activity of the immuno-captured complex (Fig. 4d).

**Fig. 4.**
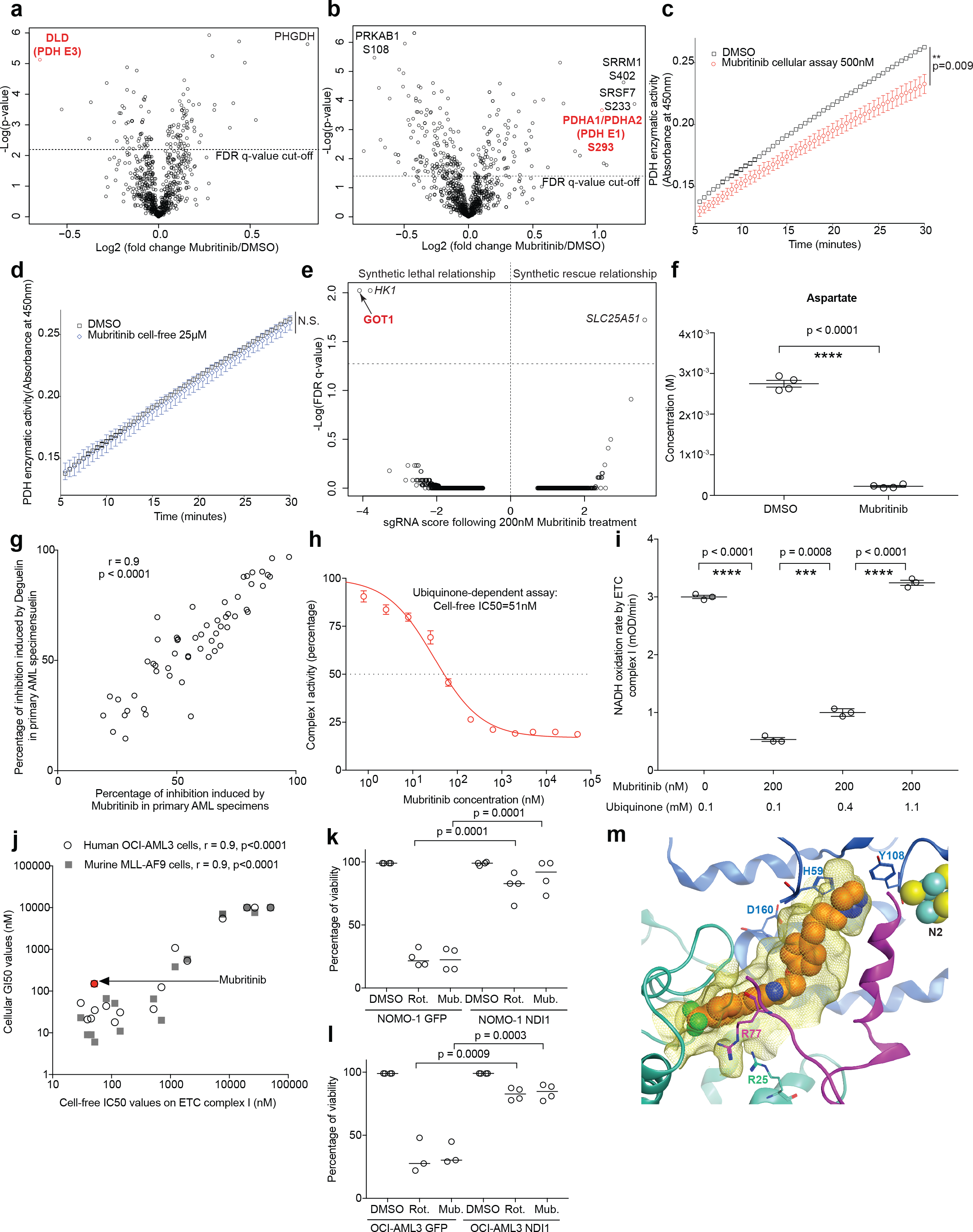
Mubritinib is a direct and ubiquinone-dependent ETC complex I inhibitor. Results of **a**, proteomic, **b**, phospho-proteomic analyses of Mubritinib treatment (500nM, 20h), in OCI-AML3 cells, average of 6 replicates. PDH enzymatic activity **c**, upon Mubritinib treatment (500nM, 20h) in OCI-AML3 cells and **d**, in a cell-free context. **e**, Results of the CRISPR/Cas9 whole genome screen in Nalm6-Cas9 clonal cells treated with 200nM Mubritinib, (see layout in Fig. S4a). **f**, Effect of Mubritinib treatment (500nM, 20h) on aspartate concentration in OCI-AML3 cells. **g**, Inhibitory patterns of Mubritinib (500nM) and Deguelin (1μM) in primary AML specimens. **h**, Mubritinib’s effect on ETC complex I activity in a cell-free ubiquinone-dependent assay (see also Fig. S4d-j). **i**, NADH dehydrogenase cell free activity in presence or absence of Mubritinib and with various concentrations of ubiquinone. **j**, Results of structure activity relationship studies in cellular and cell free assays (see also Table S8). **k**, and **l**, Rescue of cell viability through ectopic expression of yeast ETC complex I ortholog, NDI1 in NOMO-1 and OCI-AML3 cell lines, respectively after treatment with positive control Rotenone (40nM) or Mubritinib (40nM). **m**, Hypothetical binding model of Mubritinib (orange spheres) in the ubiquinone-binding pocket (yellow mesh), built in the mammalian complex (PDB entry 5LC5). Subunits 49kDa, PSST and ND1 are shown as blue, purple and green ribbons, respectively. Selected sidechains lining the site are depicted as sticks and Fe-S cluster N2 is shown as spheres. In **a-b**, statistical assessments were performed using the unpaired two-tailed *t*-test and the FDR q-value cut-off of 5% is indicated by the dotted line. In **c-d** and **f**, statistical assessments were performed using the Mann-Whitney test. In **e**, statistical assessments were performed using the Wilcoxon rank-sum test and the false discovery rate (FDR) method. In **i**, statistical assessments were performed using the two-sided unpaired *t-*test. In **j**, statistical assessments were performed using the Pearson correlation test. In **k** and **l**, statistical assessments were performed using Dunnett’s multiple comparisons test. Data in **c-d** and **f, h-i** are represented as mean values with SEM and in **k-l**, as median values.

In order to identify Mubritinib’s target pathway, we then carried out a whole-genome CRISPR/Cas9 screen in the presence or absence of Mubritinib treatment (Fig. S4a and Read files 1-3), as previously described ^44^. For this purpose, we used the B cell precursor leukemia cell line NALM-6, which, similar to poor outcome AML patient cells, is sensitive to Mubritinib (Fig. S4a), but resistant to the ERBB2 inhibitor Lapatinib (GI50>10μM, not shown). This chemo-genomic screen notably identified a synthetic lethal interaction between Mubritinib treatment and loss of *Glutamic-Oxaloacetic Transaminase 1* (*GOT1*) expression, a pyridoxal phosphate-dependent enzyme with aspartate aminotransferase activity (Fig. 4e). Accordingly, shRNA hit validations (Fig. S4b-c) revealed, among others, that silencing of *GOT1* expression in OCI-AML3 cells leads to a sensitization to Mubritinib treatment (Fig. S4b). Interestingly, recent studies reported a similar synthetic lethal interaction between *GOT1* knock-out and inhibitors of the electron transport chain (ETC). These studies showed that a major role of respiration in proliferating cells is to provide electron acceptors for aspartate synthesis, an alpha-amino acid that, in the context of ETC inhibition, can only be replenished through GOT1 activity ^45,46^. Indeed, and similar to ETC inhibitors, Mubritinib treatment in OCI-AML3 cells led to a four-fold decrease in aspartate concentrations (Fig. 4f).

So as to investigate whether Mubritinib indeed behaves as an ETC inhibitor in AML cells, we compared the inhibition pattern induced by Mubritinib treatment with that of known ETC inhibitors (Deguelin (complex I), ^47,48^ and Oligomycin (complex V)) for 56 primary AML samples (see patient characteristics in Supplementary Table S7). We found that the two ETC inhibitors’ inhibitory patterns are highly similar to that of Mubritinib (r=0.9, p<0.0001 for Deguelin, Fig. 4g and r=0.7, p<0.001 for Oligomycin, Fig. S4d). These results strongly suggest that, in the context of AML, Mubritinib targets the same molecular pathway as ETC inhibitors.

To test whether Mubritinib directly inhibits the ETC, we next carried out cell-free enzymatic activity assays for each complex of the chain. We found that, while Mubritinib did not impair the activities of ETC complexes II to V (Fig. S4e-i), it efficiently inhibited the activity of complex I (NADH dehydrogenase, *in vitro* IC50=51nM, Fig. 4h), similar to Rotenone, the reference ETC complex I inhibitor ^49^ (*in vitro* IC50=17nM, Fig. S4j). Interestingly, ubiquinone-independent diaphorase activity of NADH dehydrogenase was unaffected by Mubritinib treatment (Fig. S4k), suggesting that Mubritinib may bind at or near the ubiquinone binding site, similar to what is described for Rotenone ^50–52^. Indeed, ubiquinone supplementation was able to rescue the inhibition of NADH oxidation induced by Mubritinib treatment, demonstrating that Mubritinib acts as a ubiquinone-dependent inhibitor of NADH dehydrogenase (Fig. 4i). In addition, in agreement with an inhibitory effect on ETC complex I, Mubritinib-treated OCI-AML3 cells exhibited decreased NAD/NADH concentration ratios (Fig. S4l). NADH being the product, and NAD, the substrate of PDH, these results might explain the indirect effect of Mubritinib on PDH activity.

In order to test whether the anti-leukemic activity of Mubritinib is indeed mediated by the inhibition of ETC complex I activity, we compared the anti-leukemic potential of 15 different Mubritinib analogs in cellular assays (GI50) to their cell-free activity on ETC complex I (IC50). We found that GI50 and IC50 values strongly correlate in both human (OCI-AML3) and murine (MLL-AF9) AML cells (r=0.9, p<0.0001 for both models, Fig. 4j and Table S8). Instead of ETC complex I, *Saccharomyces cerevisiae* expresses a nucleus-encoded rotenone-insensitive NADH dehydrogenase called NDI1 ^53^, known to be able to restore OXPHOS activity in complex I deficient human cells ^54^. Accordingly, ectopic expression of NDI1 in two different human AML cell lines led to a rescue of cell viability in both Rotenone- and Mubritinib-treated cells, thus demonstrating that Mubritinib’s antileukemic activity is mainly mediated by its ability to inhibit ETC complex I activity (Fig. 4k-l).

Based on preliminary structure activity relationship studies, we propose a model of interaction between Mubritinib and the ubiquinone binding pocket of NADH dehydrogenase (Fig. 4m). Overall, our findings on Mubritinib’s mechanism of action in AML are summarized in Fig. 5a.

**Fig. 5.**
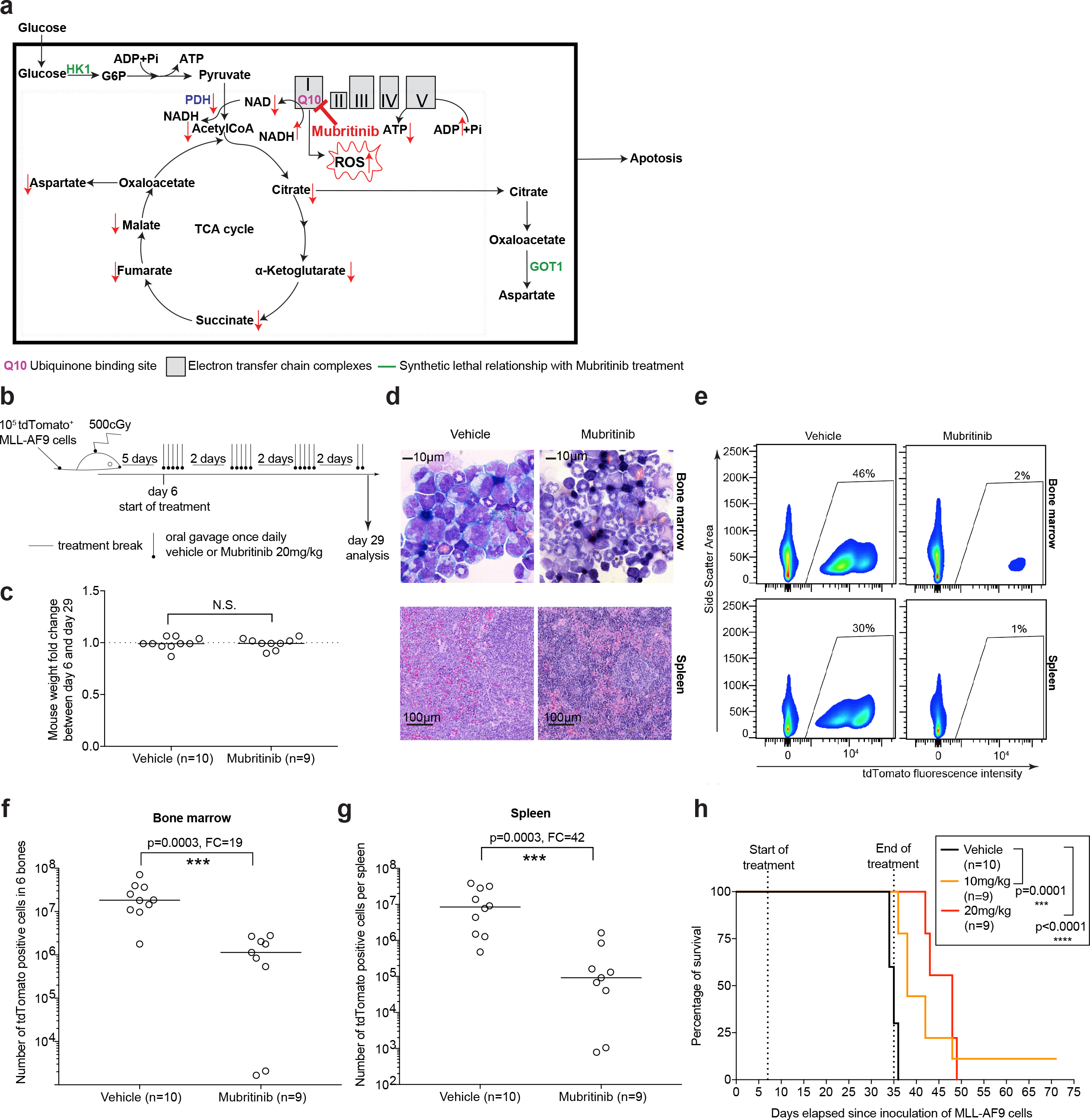
Mubritinib significantly delays the development of poor outcome AML in vivo. **a**, Summary of Mubritinib’s mode of action: briefly, Mubritinib is a direct and ubiquinone-dependent ETC complex I inhibitor. Mubritinib treatment leads to an accumulation of ROS and cells undergo apoptosis. Mubritinib treatment also induces a decrease in ATP/ADP and NAD/NADH concentration ratios, as well as in PDH, TCA and OXPHOS activities. In this context, expression of GOT1 (due to Aspartate rarefaction) and HK1 (due to OXPHOS impairment) becomes synthetic lethal. **b**, Overview of the in vivo treatment protocol. **c**, Fold change of mouse weights upon treatment. **d**, Representative bone marrow cytospins and spleen tissue sections and **e**, flow cytometry data of bone marrow and spleen hematopoietic cells at day 29 (see also Fig. S5b-c). Absolute counts of tdTomato-positive cells in **f**, the bone marrow of 6 bones (hips, femurs and tibias) and in **g**, the spleen at day 29, (see also Fig. S5d). **h**, Percentage of mice surviving from the aggressive MLL-AF9 AML model following Mubritinib treatment. FC= Fold Change. Statistical assessments in **c** and **f-g** were performed using the Mann-Whitney test and in **h** using the log-rank test. Data in **c** and **f-g** are represented as median values.

### Mubritinib significantly delays AML development*in vivo*

We next assessed the anti-leukemic potential of Mubritinib *in vivo* using the syngeneic MLL-AF9 murine AML model (Fig. 5b). MLL-AF9 cells express high levels of *HOX*-network genes and are highly sensitive to Mubritinib treatment *in vitro* (Fig. S3i) and thus represent a relevant AML model to study Mubritinib’s activity. Furthermore, the cells chosen for this study were easily trackable due to engineered expression of the fluorescent marker tdTomato. Within the treated cohort, one mouse died at day 20, probably due to compulsive gavage, as we did not detect overt leukemia development in its bone marrow (Fig. S5a). Upon analysis at day 29, mouse weights before and after treatment were not altered, both in Mubritinib and vehicle treated groups (Fig. 5c), suggesting that the treatment was overall well tolerated.

By histology, we found that the bone marrow and the spleens of treated animals contained less leukemic blasts than control animals (Fig. 5d). Accordingly, the frequencies of tdTomato positive cells in the bone marrow and the spleens of Mubritinib treated animals were largely reduced (representative plots in Fig. 5e and quantification in Fig. S5b-c). Overall, Mubritinib treatment caused a 19-fold decrease in absolute numbers of tdTomato positive cells in the bone marrow (p=0.0003, Fig. 5f) and a 42-fold decrease in the spleens (p=0.0003, Fig. 5g) of treated animals, compared to vehicle treated mice. Accordingly, Mubritinib treatment of MLL-AF9 transplanted animals significantly delayed the development of the disease in a dose dependent manner (Fig. 5h). Importantly, and in agreement with our observation that Mubritinib treatment does not affect the proliferation of normal hematopoietic CD34^+^ cells *in vitro* (Fig. 1d), the absolute number of tdTomato negative nucleated bone marrow cells was conserved after Mubritinib treatment (Fig. S5d).

In order to investigate the consequences of Mubritinib treatment on normal hematopoiesis, we treated non-transplanted PeP3B mice according to the experimental scheme in Fig. S5e. Upon analysis of the bone marrow of treated animals at day 29, we found no significant change in the absolute number of hematopoietic cells within stem or differentiated compartments (Fig. S5f-n). Analysis of blood samples at day 29 showed no significant change in neutrophil or platelet counts but revealed consistent decreases in red blood cell numbers as well as hemoglobin, hematocrit and haptoglobin measurements, all within the normal range (Fig. S5o-t). Altogether these results confirm that Mubritinib treatment does not significantly impair normal hematopoiesis.

## Discussion

Mubritinib, a small molecule developed in the early 2000s, has originally been reported as a specific receptor tyrosine kinase ERBB2 inhibitor ^35^. Mubritinib’s effect on the viability of ERBB2-negative leukemic cells led us to conclude that its mode of action in AML is likely not mediated by ERBB2 inhibition. We show that Mubritinib accumulates in the mitochondria and metabolomic investigations reveal that Mubritinib abrogates mitochondrial respiration and perturbs metabolite levels accordingly. Using proteomics, phosphoproteomics and whole genome CRISPR/Cas9 screening, as well as targeted approaches, we demonstrate that Mubritinib is in fact a novel direct and ubiquinone-dependent ETC complex I inhibitor. In line with these results, we find that the inhibitory patterns induced by Mubritinib on primary AML specimens strongly correlate with those induced by rotenoids ^11^, as well as by several ETC inhibitors, but not with those induced by the ERBB2 inhibitor Lapatinib. Interestingly, whole genome CRISPR/Cas9 screening identified loss of *Solute Carrier Family 25 Member 51* (*SLC25A51*, encoding for a mitochondrial transporter with unknown function) as having a synthetic rescue relationship towards Mubritinib treatment. Although we do not find this transporter to exhibit reduced average expression levels in Mubritinib resistant AML samples, investigation of its role in future studies might provide important information on the molecular mechanisms modulating the response to ETC complex I inhibition in leukemia. The genetic screen also revealed a synthetic lethal interaction between Mubritinib treatment and loss of *Hexokinase-1* (*HK1*), encoding an enzyme that catalyzes the first step of glucose metabolism. This result is in line with our observation that Mubritinib-treated cells switch from oxidative phosphorylation to glycolysis and is consistent with our finding that Mubritinib inhibits the ETC.

This study confirms, in the context of primary AML specimens, the recent observations from others indicating that inhibition of mitochondrial function is a promising therapeutic strategy in AML ^21,22,24,26–28^. Importantly, it also identifies and characterizes for the first time AML genetic subtypes most susceptible to respond to mitochondrial targeting. We show that, similar to normal CD34-positive cells, chemotherapy-sensitive, favorable cytogenetic risk primary AMLs do not require oxidative phosphorylation for energy production and exhibit strong transcriptomic hallmarks of hypoxia, which is consistent with recent observations made in AML xenografts ^25^. Also in line with these findings, resistance to Mubritinib associated with mutations affecting the RAS/MAPK signaling pathway, which are known inducers of glycolytic metabolism ^55,56^. In stark contrast, Mubritinib sensitivity associated with poor outcome AMLs enriched within the normal karyotype subtype, overexpressing *HOX* network genes, and carrying mutations affecting *NPM1, FLT3*-ITD as well as genes involved in the regulation of DNA methylation (*DNMT3A, TET2, IDH1,* and *IDH2*). Sensitivity in primary AML specimens also strongly correlated with increased expression of gene modules associated with mitochondrial activity, including mitochondrial respiration and indeed, direct metabolic profiling identified an association between Mubritinib sensitivity and OXPHOS hyperactivity in primary AML specimens.

In summary, using an unbiased chemo-genomic approach, our work identifies Mubritinib as a direct and ubiquinone-dependent NADH dehydrogenase inhibitor with strong *in vitro* and *in vivo* anti-leukemic activity. Importantly, we identify a genetically distinct OXPHOS-dependent population of poor outcome leukemias most susceptible to respond to ETC1-targeting, thus providing useful stratification information for the design of clinical trials testing the efficacy of OXPHOS-targeting agents. In addition, as Mubritinib already completed a phase I clinical trial in the context of ERBB2 positive solid tumors, it may rapidly and cost-sparingly be re-purposed for the treatment of such high-risk AMLs.

## Acknowledgments

The authors wish to thank Dr. Jalila Chagraoui for critical reading of the manuscript, Muriel Draoui for project coordination and Marianne Arteau and Raphaëlle Lambert at the IRIC genomics platform for RNA sequencing. We also thank the Charles-Le Moyne Hospital for providing human umbilical CB units. The dedicated work of BCLQ staff namely Giovanni d’Angelo, Claude Rondeau and Sylvie Lavallée is also acknowledged. The authors also wish to thank Jean Duchaine and the IRIC high-throughput platform staff for their helpful support for cell-based screening assays. Part of the metabolite measurement and the Seahorse analyses were performed at the Rosalind and Morris Goodman Cancer Research Centre Metabolomics Core Facility supported by the Terry Fox Foundation Oncometabolism team grant (#1048) in partnership with Fondation du cancer du Sein du Quebec (FCSQ), The Dr. John R. and Clara M. Fraser Memorial Trust, and McGill University. The authors wish to thank warmly Martin Jutras, Mihaela Friciu and Isabelle St-Jean of the Biopharmacy platform at the University of Montréal for their contribution to this study. Blood sample analysis in Fig. S5o-t was performed at the Diagnostic and Research Support Service Laboratory at the Comparative Medicine and Animal Resources Centre (CMARC), at McGill University. Finally, the authors wish to thank the authors of Molina et al. Nature Medicine 2018 for sharing plasmids for NDI1 ectopic expression. This work was mostly supported by the Government of Canada through Genome Canada and the Ministère de l’Économie, de l’Innovation et des Exportations du Québec through Génome Québec with supplementary funds from Amorchem. G.S. and J.H. are recipients of research chairs from the Canada Research Chair program and Industrielle-Alliance (Université de Montréal) respectively. BCLQ is supported by grants from the Cancer Research Network of the Fonds de recherche du Québec–Santé. RNA-Seq read mapping and transcript quantification were performed on the supercomputer Briaree from Université de Montréal, managed by Calcul Québec and Compute Canada. The operation of this supercomputer is funded by the Canada Foundation for Innovation (CFI), NanoQuébec, RMGA and the Fonds de recherche du Québec - Nature et technologies (FRQ-NT). I.Ba. has been supported by a post-doctoral fellowship from the Human Frontier Science Program (HFSP).

## Author Contributions

I.Ba. contributed to project conception, carried out most of the experiments, analyzed the results, generated the corresponding figures, tables and supplementary material and wrote the paper. Y.G. and S.G. conceived and produced the alkyne probe and the Mubritinib analogs, B.L. edited the manuscript and provided the MLL-AF9 model, J-F.S. carried out weighted gene co-expression network analyses, A.B. performed *in silico* modeling experiments, I.Bo. carried out part of the chemical screens and Seahorse analyses on primary samples, S.G. contributed to biochemical and metabolomics assays, T.MR. produced shRNAs for CRISPR/CAS9 hit validations in AML cells and edited the manuscript, J.K. carried out part of the chemical screening., NM, SC, MF and KL helped carrying out *in vivo* characterization of Mubritinib treatment, SC contributed to cell-free assay probing of analogs, C.T. carried out part of the chemical screening, VP.L. coordinated sequencing activities, E.K. carried out proteomics LC/MS experiments, T.B. supervised CRISPR/CAS9 screening experiments, J.C-H. analyzed the raw data of the CRISPR/CAS9 experiment, C.S-D. carried out the CRISPR/CAS9 experiments, G.B. is responsible for the Leucegene bioinformatics data and edited the manuscript, P.R. provided scientific guidance, M.-E.B critically commented and revised the manuscript, S.L. was responsible for supervision of the bioinformatics team and of statistical analyses., M.T. is responsible for the CRISPR/CAS9 screening activities, P.T., is responsible for the proteomics analyses, A.M. is responsible for the chemistry team as part of the Leucegene project, J.H. analyzed the cytogenetic studies, provided all the AML samples and clinical data, and edited the manuscript., and G.S. contributed to project conception and coordination and co-wrote the paper.

## Materials and correspondance

Further information and requests for resources and reagents should be directed to and will be fulfilled by the Lead Contact, Guy Sauvageau.

## Online Methods

### Experimental models and subject details

#### Primary cell cultures

This study was approved by the Research Ethics Boards of Université de Montréal, Maisonneuve-Rosemont Hospital (Montreal, QC, Canada) and Charles LeMoyne Hospital (Greenfield Park, QC, Canada). All AML samples were collected between 2001 and 2017 according to the procedures of the Banque de Cellules Leucémiques du Québec (BCLQ) and with informed consents. Detailed information about primary samples are indicated in Table S1 and Table S2. Frozen AML mono-nucleated cells were thawed at 37°C in Iscove’s modified Dulbecco’s medium (IMDM) containing 20% FBS and DNase I (100μg/mL). Cells were then cultured in optimized AML growth medium as previously reported ^6^: IMDM, 15% BIT (bovine serum albumin, insulin, transferrin; Stem Cell Technologies), 100 ng/mL SCF, 50 ng/mL FLT3-L, 20 ng/mL IL-3, 20 ng/mL G-CSF (Shenandoah), 10^−4^M β-mercaptoethanol, 500nM SR1 (Alichem), 500nM UM729 (synthesized at the Medicinal Chemistry Core Facility at the Institute for Research in Immunology and Cancer (IRIC)), gentamicin (50μg/mL) and ciprofloxacin (10μg/mL).

Control cord blood cells were collected from consenting mothers at the Charles LeMoyne Hospital (Greenfield Park, QC, Canada). Before usage in chemical screens, CD34-positive cord blood cells were isolated using the EasySep kit (StemCell Technologies, Vancouver, Canada) and cultivated for 6 days in UM171-supplemented media, as described in ^57^. During chemical screening, cord blood cells were grown in StemSpan-ACF (Stemcell Technologies 09855) containing SCF 100ng/mL, TPO 50ng/mL, FLT3-L 100ng/mL, Glutamax 1X, LDL 10μg/mL and ciprofloxacin (10μg/mL) as well as 500nM SR1 and 35nM UM171.

#### Cell lines

BT474 female cells were a kind gift from the laboratory of Sylvie Mader and were cultured in DMEM 10% FBS. OCI-AML2, OCI-AML3 and OCI-AML5 male cells were provided by the The University Health Network (Toronto). OCI-AML5 cells were expanded in alpha-MEM, 20% heat-inactivated FBS, 10ng/mL GM-CSF (Shenandoah). OCI-AML2 and OCI-AML3 cells were cultured in alpha-MEM, 20% heat-inactivated FBS. NOMO-1 female and NALM-6 male cells were purchased from DSMZ and cultured in RPMI 1640, 10% heat-inactivated FBS. HEK-293T female cells were purchased from ATCC and grown in alpha-MEM 10% FBS.

Murine leukemias (female cells) were generated by infection with VSV-G-pseudotyped MSCV MLL-AF9 IRES Puro (subcloned from a construct by Frédéric Barabé, Laval U, Québec, QC, Canada) as described before ^7^. Secondary infections of MLL-AF9 cells with tdTomato expressing MSCV vectors were done by GPE+86 co-culture under the same conditions or by spinoculation (1,000g, 32°C, 2h) with VSV-G pseudotyped Plat-A virus-containing supernatant in the presence of polybrene. Murines were cultivated in (IMDM, 10% heat-inactivated FBS, 100 ng/mL rmSCF (Shenandoah), 10 ng/mL rmIL-3 (Shenandoah), 10 ng/mL rhIL-6 (Shenandoah), 10^-4^M 2-mercaptoethanol.

All cell lines were grown in humidified incubators at 37 degrees Celsius and 5% CO_2_.

#### Animals

All animal procedures complied with recommendations of the Canadian Council on Animal Care and were approved by the Deontology Committee on Animal Experimentation at University of Montreal. For the MLL-AF9 study, C57BL/6J female mice were purchased from Jackson Laboratory (Bar Harbor, Maine, #000664) and bred in a pathogen-free animal facility. 8 to 12-week-old sub-lethally irradiated (500cGy, ^137^Cs-gamma source) C57BL/6J mice were used. For the study of Mubritinib’s effect on normal hematopoiesis *in vivo*, B6.SJL-*Ptprc*^*a*^ *Pepc*^*b*^/BoyJ (Pep3B) female mice were purchased from Jackson Laboratory (Bar Harbor, Maine, #002014) and bred in a pathogen-free animal facility. 8 to 12-week-old sub-lethally irradiated (500cGy, ^137^Cs-gamma source) Pep3B mice were used. For all experiments, littermates of the same sex were randomly assigned to experimental groups.

### Method details

#### Chemical screens

All powders were dissolved in DMSO and diluted in culture medium immediately before use. Final DMSO concentration in all conditions was 0.1%. Cells were seeded in 384-well plates in 50μL media per well at the following densities: AML patient cells, 5,000 cells per well; Cord blood cells, 2,000 cells per well; OCI-AML3 cells, 150 cells per well; BT474 cells, 2000 cells per well; MLL-AF9 cells and HOXA9/MEIS1 cells, 90 cells per well; AML/ETO cells, 1350 cells per well. Compounds were added to seeded cells in serial dilutions (10 dilutions, 1:3 or 8 dilutions, 1:4), in duplicates or quadruplicates. Cells treated with 0.1% DMSO without additional compound were used as negative controls. Viable cell counts per well were evaluated after 5 days of culture (for murine cells) or 6 days of culture (for human cells) using the CellTiter-Glo assay (Promega) according to the manufacturer’s instruction. The percent of inhibition was calculated as follows: 100-(100×(mean luminescence(compound)/mean luminescence(DMSO)); where mean-luminescence(compound) corresponds to the average of luminescent signals obtained for the compound-treated cells, and mean-luminescence(DMSO) corresponds to the average of luminescent signals obtained for the control DMSO-treated cells.

#### Assessment of Mubritinib’s activity *in vivo*

MLL-AF9 cells used in this study originate from the diseased bone marrow of a primary recipient C57BL/6J female mouse transplanted with MSCV MLL-AF9 ires Puro-T2A-rtTA2 and MSCV ires tdTomato infected cells as described in ^7^. Briefly 100,000 such cells were transplanted *via* the tail vein into 8 to 12-week-old sub-lethally irradiated (500cGy, ^137^Cs-gamma source) C57BL/6J mice. In a first experiment, mice were fed once daily by oral gavage either vehicle (0.86% Natrosol/14% DMSO solution, 10μL/g of mouse, n=10 recipient mice) or 20mg/kg Mubritinib (n=9 recipient mice), following the experimental scheme shown in Fig. 5a. Mice were sacrificed at day 29 and their bone marrow (2 hips, 2 femurs and 2 tibias) and spleens were analyzed by histology (bone marrow cytospins were colored using the Wright Giemsa stain and Hematoxylin and Eosin staining was used for paraffin sections of spleens), as well as by flow cytometry. In a second experiment, we counted the number of mice dying from leukemia after being fed once daily by oral gavage either vehicle (n=10 recipient mice) 10mg/kg or 20mg/kg Mubritinib (n=9 recipient mice each), following the experimental scheme shown in Fig. 5a, extended until day 35. Mice were sacrificed when they showed marked leukemic symptoms. AML development in sacrificed animals was confirmed by detection of high percentages (>80%) of tdTomato positive cells in the bone marrow and spleens by flow cytometry. For the evaluation of Mubritinib’s effect on normal hematopoiesis, 8 to 12-week-old sub-lethally irradiated (500cGy, ^137^Cs-gamma source) Pep3B mice were fed once daily by oral gavage either vehicle (0.86% Natrosol/14% DMSO solution, 10μL/g of mouse, n=5 recipient mice) or 10mg/kg Mubritinib (n=5 recipient mice), following the experimental scheme shown in Fig. S5e. Mice were sacrificed at day 29 and bone marrow (analysis by flow cytometry, see below and Key resources Table) as well as blood samples were collected. Cell blood counts were performed at the McGill Diagnostic and Research Support Service Laboratory at the Comparative Medicine and Animal Resources Centre (CMARC), using the scil Vet ABC Plus hematology analyzer: WBCs, RBCs, and platelets were counted via electrical impedance, hemoglobin was measured by photometry, and hematocrit was calculated. Haptoglobin levels were measured by photometric methodology by IDEXX Laboratories (Markham, Ontario), using a Cobas 6000 automated chemistry analyzer.

#### Flow cytometry analyses

Antibodies used in the study are listed in the Key Resources Table. Dead cells were stained using Propidium iodide at a final concentration of 1μg/mL. For reactive oxygen species quantification, cells were stained with 1μM H2DCFDA (Thermo Fisher, D399) for 30 minutes under normal growth conditions. For absolute cell counts, counting beads (CountBrightTM Absolute Counting Beads by Molecular Probes) were added to FACS tubes. Cells were analyzed on LSRII flow cytometer (BD Bioscience), BD Canto II cytometer (BD Bioscience) or on an IQue Screener (Intellicyt) and results were analyzed with BD FACS Diva 4.1 and FlowJo softwares.

#### Next-generation sequencing and mutation quantification

Workflow for sequencing, mutation analysis and transcripts quantification of the Leucegene cohort have been described previously ^12^. Briefly, libraries were constructed with TruSeq RNA / TruSeq DNA Sample Preparation Kits (Illumina). Sequencing was performed using an Illumina HiSeq 2000 with 200 cycles paired end runs. Sequence data were mapped to the reference genome hg19 using the Illumina Casava 1.8.2 package and Elandv2 mapping software according to RefSeq annotations (UCSC, April 16^th^ 2014). Variants were identified using Casava 1.8.2 and fusions or larger mutations such as partial tandem duplications with Tophat 2.0.7 and Cufflinks 2.1.1. Transcript levels are given as Reads Per Kilobase per Million mapped reads (RPKM) and genes are annotated according to RefSeq annotations (UCSC, April 16th 2014).

#### Weighted gene co-expression network analysis (WGCNA)

Sequencing of the 200 primary specimens was performed using the Illumina HiSeq 2000 device with 200 cycles paired-end runs. Resulting reads were aligned to the Genome Reference Consortium Human Build 38 patch release 84 (GRCh38.84) using STAR aligner v.2.5.1 ^58^ and counted with the RNA-Seq by Expectation Maximization (RSEM) software v1.2.28 ^59^.

The R package WGCNA ^38^ was used to perform a weighted correlation network analysis with normalized expression data (TPM) as input. Co-expression similarities were obtained by calculating Pearson’s correlations between genes. Adjacencies were computed by raising co-expression similarities to a power β = 12 (soft thresholding). β was chosen as the lowest integer allowing the resulting network to exhibit an approximate scale-free topology (as advised in the original method ^38^). The adjacency matrix obtained was transformed into Topological Overlap Matrix (TOM) ^60^ and a clustering tree constructed by average linkage hierarchical clustering using the corresponding dissimilarities (1-TOM). Module detection was then conducted using the Dynamic Tree Cut method ^61^ (minimum module size = 20; deepSplit = 3). Correlations between eigengenes (first principal component of each module) and Mubritinib GI50 values were computed and significance assigned to each association. A Gene Significance (GS) was also assigned to each individual gene composing a module. A GO terms enrichment analysis using the hyperGTest function (ontology = ‘BP’, pvalueCutoff = 0.05; testDirection = ‘over’) from the GOstats package ^62^ was performed for each module using significantly associated genes (p.GS < 0.05).

#### LC/MS metabolite measurements

NAD, NADH, ATP, ADP, AMP and Acetyl CoA measurements were carried out at the Biopharmacy platform at Université de Montréal. OCI-AML3 cells (treated with either DMSO or Mubritinib 500nM for 20h) were washed with ice cold 150mM NH4 formate (pH 7.4) and resulting cell pellets lysed by adding 400μL of ice-cold 65% (v/v) methanol / 50mM NH4HCO3, pH 8,0 on dry ice, followed by 2 min of sonication in a sonic water bath. After a 20 min incubation period on dry ice, lysates were centrifuged at 14,000g for 5 min at 4°C to pellet the cell debris. The polar metabolite-containing supernatants were transferred to new tubes on dry ice and kept at −80°C until analysis. Quantification of metabolites was performed using a triple quadrupole mass spectrometer (SCIX API4000 QTRAP) and the metabolites were separated using a pH gradient on a weak anion exchange column (Thermo Biobasic AX) in HILIC mode.

All other LC/MS metabolite measurements were carried out at the McGill Metabolomics core facility. Authentic metabolite standards were purchased from Sigma-Aldrich Co., while the following LC/MS grade solvents and additives were purchased from Fisher: ammonium acetate, formic acid, water, methanol, and acetonitrile. OCI-AML3 cells (5 million cells, quadruplicates, treated with either DMSO or Mubritinib 500nM for 20h) were washed twice with ice-cold 150mM ammonium formate pH 7.2. Metabolites were then extracted using 380μl of LC/MS grade 50% methanol/50% water mixture and 220μl of cold acetonitrile. Samples were then homogenized by the addition of 6 1.4mm ceramic beads and bead beating 2min at 30Hz (TissueLyser, Qiagen). A volume of 300μl of ice-cold water and 600μl of ice-cold methylene chloride were added to the lysates. Samples were vortexed and allowed to rest on ice for 10 min for phase separation followed by centrifugation at 4,000rpm for 5min. The upper aqueous layer was transferred to a fresh pre-chilled tube. Samples were dried by vacuum centrifugation operating at -4°C (Labconco) and stored at −80°C until ready for LC-MS/MS data collection.

For targeted semi-quantitative analysis of Amino acids or nucleotides LC-MS/MS was utilized. Specimens were first re-suspended in 50μL of water and clarified by centrifugation for 5 min at 15,000 rpm at 1°C. Samples were maintained at 4°C for the duration of the LC-MS/MS analysis in the autosampler. Analysis of nucleotides was performed first followed by analysis of amino acids and citric acid cycle intermediates. Multiple reaction monitoring (MRM) transitions were optimized on standards for each metabolite analyzed. Data were quantified by integrating the area under the curve of each sample compound using MassHunter Quant (Agilent Technologies) and compared to a dilution series of authentic standard dissolved in water. These data are considered semi-quantitative due to potential uncorrected matrix effects.

For nucleotide analysis, a volume of 5μL was injected an Agilent 6430 Triple Quadrupole (QQQ) equipped with a1290 Infinity ultra-performance LC system (Agilent Technologies, Santa Clara, CA, USA). Separation was achieved using Scherzo SM-C18 column 3μm, 3.0x150mm (Imtakt Corp, JAPAN) maintained at 10°C. The chromatographic gradient started at 100% solvent A (5mM ammonium acetate in water) followed by a 5min gradient to 100 % solvent B (200mM ammonium acetate in 20% acetonitrile). The gradient was held at 100% solvent B for 5 min. The flow rate was set for 0.4mL/min. The column was then re-equilibrated at 100% solvent A for 6 min before the next injection.

After extensive re-equilibration to a different solvent system, the same column and instrument were used for amino acid and citric acid cycle intermediate detection. Separation was achieved using a gradient starting gradient started at 100% mobile phase A (0.2% formic acid in water) with a 2min hold followed with a 6min gradient to 80% B (0.2 % formic acid in MeOH) at a flow rate of 0.4 ml/min and column temperature of 10°C. This was followed by a 5min hold time at 100% mobile phase B and a subsequent re-equilibration time (6 min) before the next injection.

### LC/MS proteomics and phospho-proteomics approaches

#### Protein extraction and enzymatic digestion

Cells were lysed by sonication in lysis buffer (1% sodium deoxycholate, 100 mM NH_4_HCO_3_ pH 8.0, 10 mM TCEP and 40 mM chloroacetamide supplemented with HALT phosphatase inhibitor cocktail, Pierce) with subsequent 5 min incubation at 95°C. Samples were centrifuged at 40,000 x g for 10 min, and the supernatants were transferred into clean tubes prior to determination of protein concentrations by BCA-RC assay (Thermo Fisher Scientific). Proteins were digested with trypsin (Sigma-Aldrich) overnight at 37°C using an enzyme to substrate ratio of 1:50 (w/w). Tryptic digests were acidified with FA to final of 1% (v/v), centrifuged (20,000 x g 10 min) and desalted on Oasis HLB cartridges (Waters) according to manufacturer instructions. Peptide eluates were snap-frozen in liquid nitrogen, lyophilized in a speedvac centrifuge and stored at −80°C.

#### Phosphopeptide isolation and fractionation

Tryptic digests were subjected to enrichment on TiO_2_ beads as described previously ^63^. Sample loading, washing, and elution steps were performed using custom StageTips ^64,65^ made from 200 μL pipette tips containing a SDB-XC membrane (3M) frit and filled with TiO_2_ beads. We equilibrate TiO_2_ material in 250 mM lactic acid 70% ACN 3% TFA, the same buffer is used for sample loading. After extensive wahing steps retained phosphopeptides were displaced from TiO_2_ with 500 mM phosphate buffer at pH=7. Peptides were desalted in 50 μL of 1% FA directly on SDB-XC frits and subsequently eluted using 50 μL of 50% acetonitrile (ACN) 1% FA. Eluates were dried in a speedvac and stored at −80°C. Prior to LC-MS/MS analyses peptides were resuspended in 10 μL of 4%.

#### Mass spectrometry

Peptides were analyzed by LC-MS/MS using a Proxeon nanoflow HPLC system coupled to a tribrid Fusion mass spectrometer (Thermo Fisher Scientific). Each sample was loaded and separated on a reverse-phase analytical column (18 cm length, 150 μm i.d.) (Jupiter C_18_, 3μm, 300 Å, Phenomenex) packed manually. LC separations were performed at a flow rate of 0.6 μL/min using a linear gradient of 5-30 % aqueous ACN (0.2% FA) in 106 minutes. MS spectra were acquired with a resolution of 60,000. “TopSpeed” (maximum number of sequencing events within 5 sec window) method was used for data dependent scans on the most intense ions using high energy dissociation (HCD). AGC target values for MS and MS/MS scans were set to 5e5 (max fill time 200 ms) and 5e4 (max fill time 200 ms), respectively. The precursor isolation window was set to m/z 1.6 with a HCD normalized collision energy of 25. The dynamic exclusion window was set to 30s.

#### Data processing and analysis

MS data were analyzed using MaxQuant^66,67^ software version 1.3.0.3 and searched against the SwissProt subset of the *H. Sapiens* uniprot database (http://www.uniprot.org/). A list of 248 common laboratory contaminants included in MaxQuant was also added to the database as well as reversed versions of all sequences. The enzyme specificity was set to trypsin with a maximum number of missed cleavages set to 2. Peptide identification was performed with an allowed initial precursor mass deviation up to 7 ppm and an allowed fragment mass deviation of 20 ppm with subsequent non-linear mass re-calibration^68^. Phosphorylation of serine, threonine and tyrosine residues was searched as variable modification; carbamidomethylation of cysteines was searched as a fixed modification. The false discovery rate (FDR) for peptide, protein, and site identification was set to 1% and was calculated using decoy database approach. The minimum peptide length was set to 6, and the ‘peptide requantification’ function was enabled. The option match between runs (1 min time tolerance) was enabled to correlate identification and quantitation results across different runs. In addition to an FDR of 1% set for peptide, protein and phosphosite identification levels, we considered only phosphosites for which localization confidence was higher than 75%. Relative quantification of the peptides against their heavy-labeled counterparts was performed with MaxQuant using area under 3D pepride peak shapes^66,67^.

#### Whole genome CRISPR/Cas9 screen

The Extended Knockout (EKO) pooled lentiviral library of 278,754 sgRNAs targeting 19,084 RefSeq genes, 3,872 hypothetical ORFs and 20,852 alternatively spliced isoforms was introduced within a clone of the NALM-6 pre-B lymphocytic cells line with a doxycycline-inducible Cas9 was described previously 44. NALM-6 cells at 200,000 cells per ml were exposed for a period of 3 days to various concentrations of Mubritinib and luminescence output following addition of the cell viability CellTiter-Glo assay reagent (Promega) was measured with a Biotek Synergy Neo multi-mode microplate reader. With this pattern of NALM-6 growth inhibition in response to amount of compound used (GI50 value close to 500 nM), we estimated that 200nM would inhibit growth sufficiently to observe growth rescue phenotypes while still allowing enough growth to observe drug sensitivity phenotypes. The EKO library (kept at a minimum of 250 cells per sgRNA) was thawed and cultured in 10% FBS RPMI supplemented with 2 ug/mL doxycycline for a period of 7 days to induce knockouts with dilutions to 400,000 cells per ml every 2 days. At day 7, 70 million cells were spun at 1,200 rpm for 5 min, washed with 1X PBS, pelleted and frozen (Day 7 control). The library was left to expand 8 more days without doxycycline with 200 nM Mubritinib (a total of 100 cells per sgRNA on average) or DMSO only (250 cells per sgRNA). Cell concentration was assessed every 2 days and cells diluted back to 400,000 cells per ml whenever cell concentration was higher than 800,000 cells per ml. During this period, there were 7.6 population doublings for the DMSO control while the treated cells had only 3.5. Both samples were then PBS-washed and cell pellets frozen. Genomic DNA was extracted from all 3 samples using the QIAamp DNA blood maxi kit (Qiagen). SgRNA sequences were recovered and fitted with Illumina adaptors by PCR and NGS performed on an Illumina HiSeq 2000 device (Génome Québec Innovation Center) as previously described ^44^. Synthetic rescue/positive selection scores were determined using the RANKS algorithm ^44^ using sgRNA read numbers of the treated sample compared to the Day 7 control whereas synthetic lethal/negative selection scores were calculated by comparing the treated sample to the Day 15 control.

#### Molecular modelling

The MOE package (Molecular Operating Environment (MOE), 2018.0101; Chemical Computing Group ULC, 1010 Sherbooke St. West, Suite #910, Montreal, QC, Canada, H3A 2R7, 2018) was used for structural analysis, protein protonation (Protonate 3D approach, Amber10:EHT forcefield), molecular docking and rendering. Docking of Mubritinib and analogs was performed on the bovine complex I structure (PDB entry 5LC5), ^69^ and the search area was restricted to the ubiquinone binding site. The theoretical binding model of Mubritinib was derived from the analysis of the best docking poses obtained with Mubritinib and the active analogs.

#### Click chemistry

BT474 for were plated on IBIDI 6-well slides the day prior to the experiment at a density of 200,000 cells per well. The next day, cells were treated for 2 hours with the alkyne Mubritinib probe at 10μM in normal growth conditions. Cells were then stained with Mitotracker green 100nM for 15min (ThermoFisher Scientific, M7514), or ERBB2-FITC 1:10 for 20min (Biolegend, 324404). Cells were then washed twice in PBS, 2%FBS and the click reaction was carried out in a solution of copper (2.5mM) and ascorbate (5mM) in PBS together with a Cy5 click dye (1:200) at room temperature for one hour. Cells were then washed in PBS and counterstained with DAPI (1:2000) for 2min. Pictures were taken on a confocal microscope LSM700, using the Software ZEN lite from ZEISS Microscopy.

#### shRNA validations

Lentiviral vectors carrying shRNAs targeting candidate genes were generated by cloning appropriate shRNA sequences as described in ^70^ into the MNDU-GFP-miRE vector. Control vector (shLUCI.1309) contained shRNA targeting luciferase.

HEK293T cells were transfected with 0.75μg lentiviral plasmid, 0.5μg PAX2 packaging plasmid and 0.15μg VSV-G envelope plasmid using 3μL JetPrime Transfection reagent (PolyPlus Transfection), according to manufacturer’s directions. Viral supernatant was collected after 48 hours, filtered and 0.5mL was added fresh to 300,000 OCI-AML3 cells in a 24 well plate, with the addition of 10μg/mL protamine sulfate. Cells were infected by spinoculation at 1,000g for 2hrs at 32°C. Virus was washed off cells after overnight incubation and cells re-plated in alpha-MEM 20% FBS. Infection efficiency (%GFP positive) and cell counts were assessed 48hr post-infection using an iQue Flow cytometer (Intellicyt) and Tru-count beads (BD Bioscience). Infection of OCI-AML3 cells resulted in 70–98% GFP+ cells. Knockdown assessments were carried out 2 to 4 days post-infection: RNA was harvested in Trizol (ThermoFisher) and isolated according to manufacturer’s protocol and reverse transcribed using MMLV reverse transcriptase and random primers (ThermoFisher). Quantitative PCR was performed for shRNA target genes using validated assays designed for the Universal Probe Library (Roche) on the Viia7 (Applied Biosystems). Relative quantity of target is normalized to HPRT and compared to normalized expression in shLUCI.1309 infected control cells.

#### NDI1 ectopic expression

HEK293T cells were transfected with 0.75μg lentiviral plasmid (pLenti6.3/V5 NDI-1 or pLenti6.3/V5 GFP), 0.5mg PAX2 packaging plasmid and 0.15μg VSV-G envelope plasmid using 3μL JetPrime Transfection reagent (PolyPlus Transfection), according to manufacturer’s directions. Viral supernatant was collected after 48 hours, filtered and fresh viral supernatant (0.5mL) was added to 300,000 OCI-AML3 cells in a 24 well plate, in the presence of 10mg/mL protamine sulfate. Cells were infected by spinoculation at 1,000g for 2hrs at 32°C. Virus was washed off cells after overnight incubation and cells re-plated in alpha-MEM 20% FBS. After infection, transduced cells were selected through growth in 7 μg/mL blasticidin. Blasticidin selected cells were subsequently treated for 6 days with either DMSO, Rotenone (40nM) or Mubritinib (40nM). Cell viability was measured using CellTiter-Glo (Promega).

#### Leukemic stem cell (LSC) frequency assessment

LSC frequencies were assessed in immunocompromised NSG mice using limiting dilution assays, as detailed previously ^10^. NOD.Cg-Prkdc^*scid*^ Il2rg^*tm1Wjl*^/SzJ (NSG) mice were purchased from Jackson Laboratory (Bar Harbor, Maine) and bred in a pathogen-free animal facility. AML samples were transplanted via the tail vein into 8 to 12-week-old sub-lethally irradiated (250cGy, ^137^Cs-gamma source) NSG mice. AML cells were transplanted at four different cell doses in groups of four recipient mice directly after thawing. Human leukemic engraftment in mouse bone marrow was determined by flow cytometry between 10- and 16-weeks post-transplant. On average 150,000 gated events were acquired. Mice were considered positive if human cells represented > 1% of the bone marrow cell population. Mice were excluded only in case of obvious non-leukemia related death (*e.g*., first two weeks after irradiation). To discriminate between engraftment of leukemic and normal cells present in unsorted patient samples only recipients with proportions of CD45^+^CD33^+^ or CD45^+^CD34^+^ cells higher than proportions of CD19^+^CD33^-^ or CD3^+^ were considered to harbor cells of leukemic origin.

#### Enzymatic activity assays

PDH activity kit was purchased from Abcam, Cambridge, UK (ab109902) and the assay was carried out following the manufacturer’s recommendations using proteins extracted from OCI-AML3 cells exposed to DMSO or Mubritinib 500nM for 20h, or proteins extracted from OCI-AML3 cells exposed to Mubritinib (25μM) after immunocapture of the complex.

The cell-free activity assays for the different ETC complexes were purchased from Abcam: complex I: ab109903, complex II+III: ab109905, complex IV: ab109906, complex V: ab109907. All kits were used in accordance with the manufacturer’s protocol. Cell-free IC50 values (concentration inducing 50% of inhibition of the enzyme’s activity) were calculated using GraphPad^®^ Prism 4.03 (La Jolla, CA, USA) by four-parameter-non-linear curve fitting methods. Finally, the ubiquinone-independent diaphorase complex I activity kit was purchased from Abcam (ab109721) and the assay was carried out following the manufacturer’s recommendations using proteins extracted from OCI-AML3 cells exposed to DMSO or Mubritinib 500nM for 20h.

#### Seahorse metabolic flux experiments

Oxygen consumption rates and extracellular acidification rates were measured using a 96-well Seahorse Bioanalyzer XFe96 or XFe24 according to the manufacturer’s instructions (Agilent Technologies). Seahorse XF Base medium was supplemented with 1mM pyruvate, 2mM glutamine and 10mM glucose in the case of Mitochondrial Stress Test and with 1mM pyruvate, 2mM glutamine and no glucose in the case of Glycolytic Stress Test. The pH of the Seahorse media was then adjusted at 7.4 prior to assay. In brief, leukemic cells were seeded into Seahorse 96-well (or 24-well) plates pre-coated for 3 h with poly-lysine (Sigma-Aldrich, P4707) at a density of 125,000 cells/well in 100uL (or 150,000 cells/well, in 150μL) of temperature/CO_2_ pre-adjusted Seahorse media per well. The Seahorse plates were then centrifuged at 1400rpm for 5 min. An additional 75μL (or 375uL) of Seahorse media was then added and cells were eventually analyzed following the manufacturer’s instructions by adding compounds in a constant volume of 25μL (or 75μL). Compounds were acutely injected in cells at a final concentration of 1μM for Mubritinib, 1μM for Oligomycin, 0.5μM for FCCP, 0.5μM for Rotenone/Antimycin A, Glucose 10mM and 2-Deoxy-Glucose 50mM. Leukemic cells from cell lines were passaged in fresh standard culture media 24 prior to being harvested for Seahorse analysis. Leukemic cells from primary AML specimens were thawed and cultured for 24h as described in “Primary cell cultures” section before being harvested for Seahorse analysis.

### Quantification and statistical analysis

Analysis of differential gene expression was performed using the Wilcoxon rank-sum test and the false discovery rate (FDR) method was applied for global gene analysis as previously described ^12^. Differential overall-survival p-values were calculated using the log-rank test. GI50 values (corresponding to the concentration of compound required to reach 50% of inhibition) and cell free IC50 values were calculated using ActivityBase SARview Suite (IDBS, London, UK) and GraphPad Prism 4.03 (La Jolla, CA, USA) by four-parameter-non-linear curve fitting methods. Statistical probing methods for each figure are indicated in the corresponding figure legends.

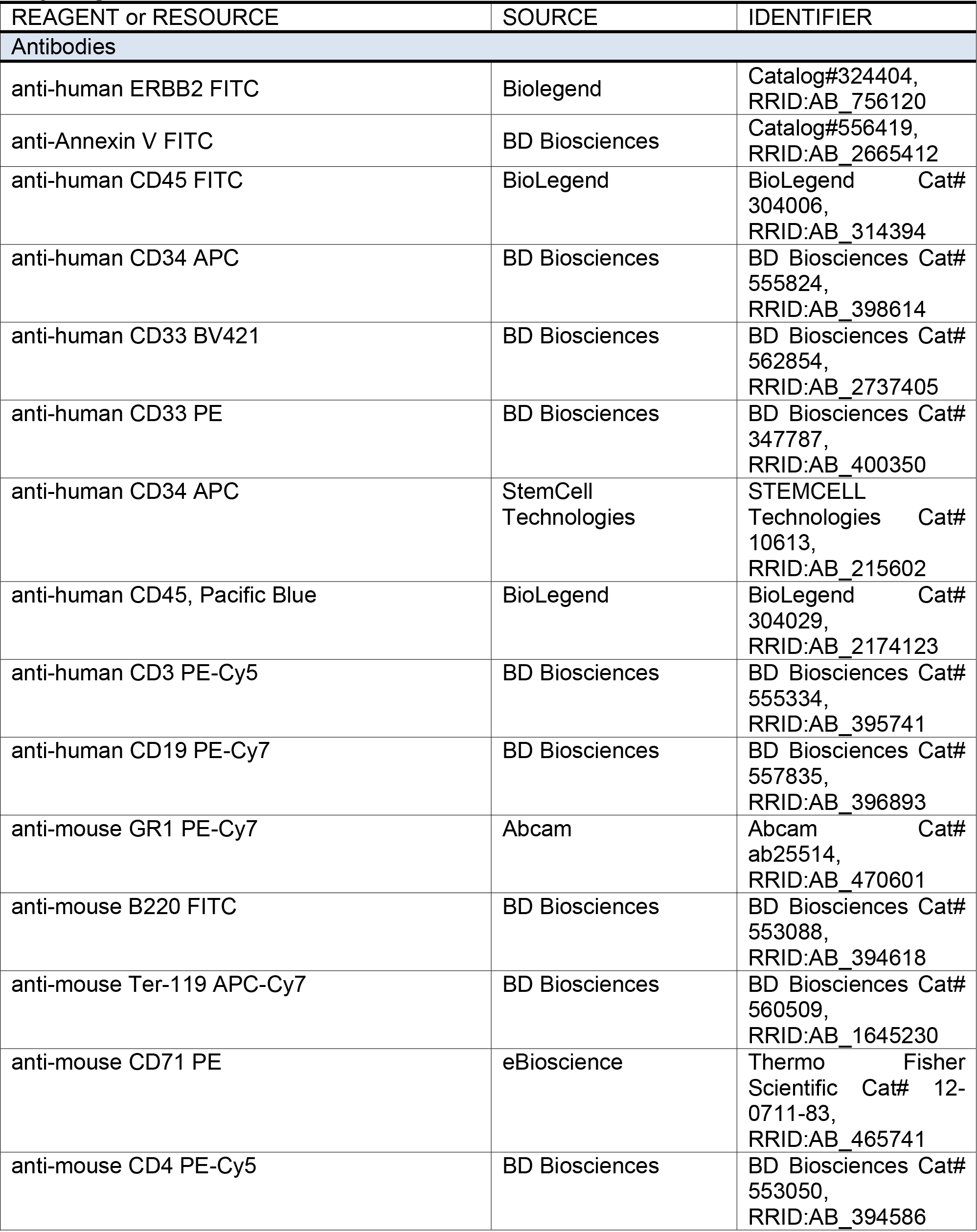
**Key Reagents And Resources Table**

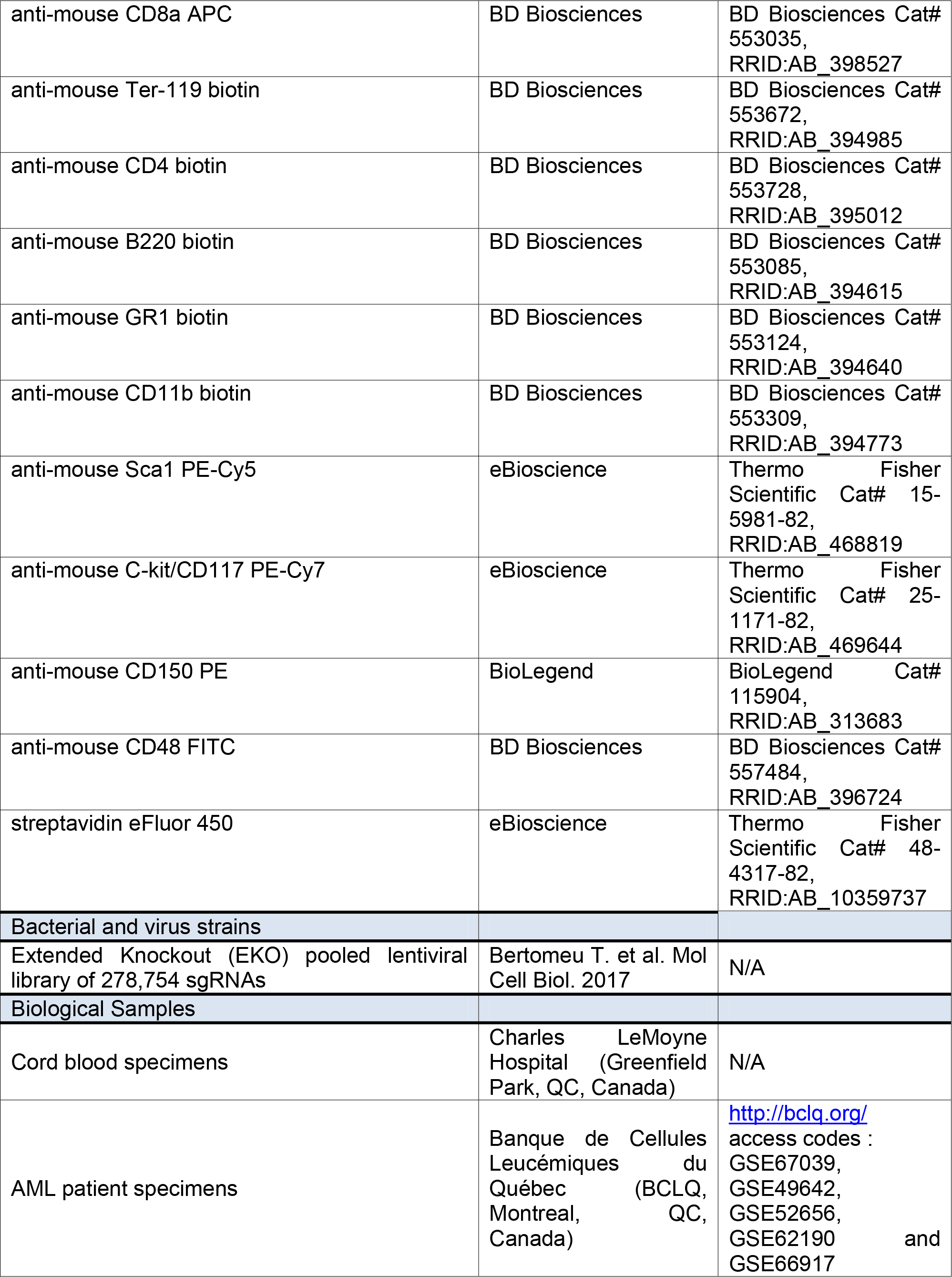

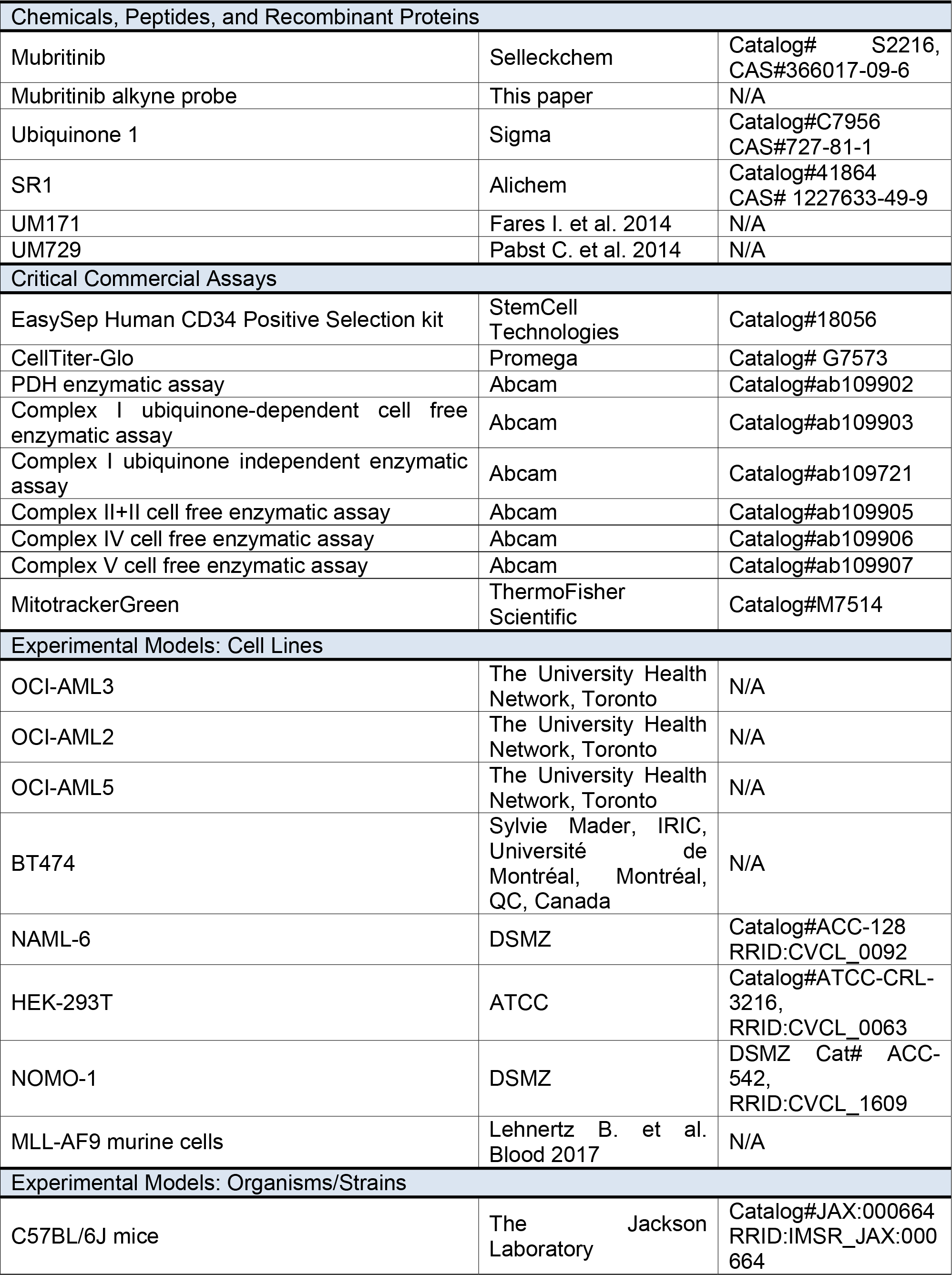

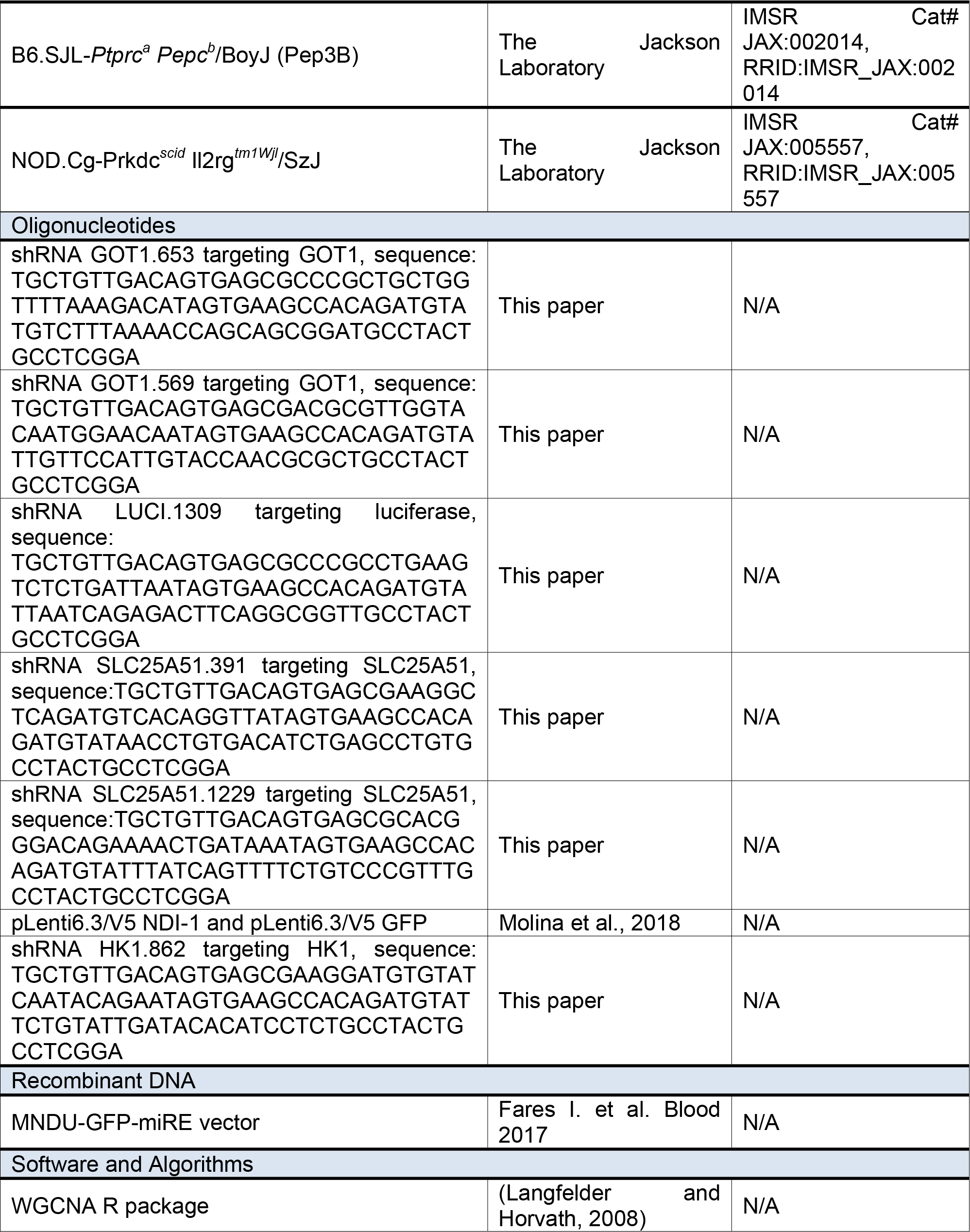

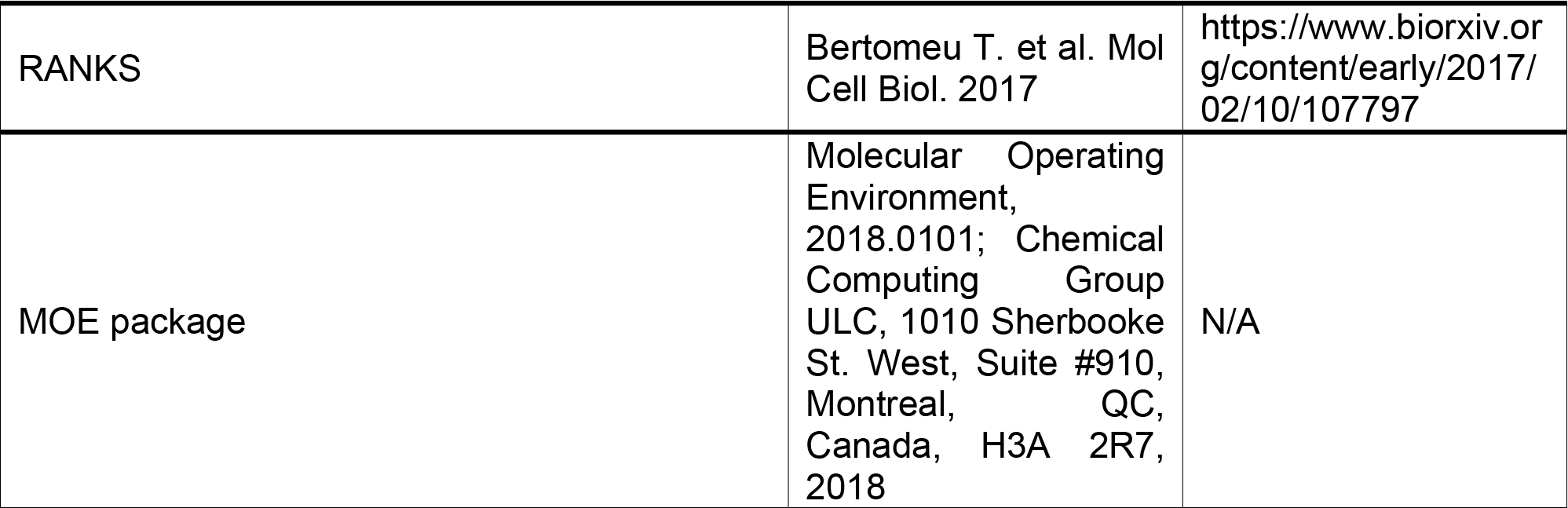

