## Supplementary Information PDF for "Genetic Landscape of Electron Transport Chain Complex I Dependency in Acute Myeloid Leukemia"

### Supplemental Figure and Table Legends

**Fig. S1, related to Fig. 1: Clinical, mutational and transcriptional landscape of poor outcome AML specimens.**

**a.** Frequency of poor outcome AMLs according to their cytogenetic risk classification. **b.** Clinical features and **c.** mutations enriched in poor *versus* good outcome AML specimens according to a Bonferroni-corrected exact Fisher's test. Significant enrichments are displayed in bold red when associated with poor outcome and in bold blue when associated with good outcome AMLs. **d.** Transcriptomic profiles of poor *versus* good outcome specimens, highlighting differentially expressed genes with average expression levels strictly above 0.1RPKM and with differences in average expression strictly above 10-fold. Analysis of differential gene expression in **d.** was performed using the Wilcoxon rank-sum test and the false discovery rate (FDR) method.

**Fig. S2, related to Fig. 2: Clinical and mutational landscape of Mubritinib sensitive AML specimens.**

**a.** *In vitro* fold-expansion of Mubritinib-resistant (GI50≥374nM, n=100) and –sensitive (GI50<374nM, n=100) cells in SR1 + UM729 (S7) conditions [6] **b.** Mubritinib GI50 values in triple mutated AMLs compared to all other tested specimens. Statistical assessments were performed using the Mann-Whitney test and data are represented as median values.

**Fig. S3, related to Fig. 3: Dissection of Mubritinib's mechanism of action.**

**a**. ERBB2 gene expression in Mubritinib-sensitive *versus* Mubritinib-resistant patient samples as compared to positive control breast tumor or healthy tissues. **b.** Dose response assay of OCI-AML3 and OCI-AML5 human AML cell lines representing human models of Mubritinib-sensitive *versus* -resistant AML. **c.** Structure of the Mubritinib alkyne probe used for click-chemistry experiments and its signal (Cy5) detected by flow cytometry with or without competitive co-treatment with Mubritinib as indicated. **d.** Control and **e.** Mubritinib-induced (1uM acute injection, as indicated) measurements of extracellular acidification rates in two human AML cell lines using a Seahorse analyzer. Intracellular **f.** and extracellular **g.** lactate concentrations in CD34-positive cord blood cultures after Mubritinib treatment (500nM, 20h). **h.** Dose response assay of MLL-AF9 (2 independent infections) murine models of AML. **i**. Apoptotic cell death rates in MLL-AF9 cells upon Mubritinib treatment (500nM), (see also Fig. 3p). **j.** Oxygen consumption rate profiles of Mubritinib sensitive and resistant primary AML specimens (see cohort in Table S6). **k.** Basal extracellular acidification rates measured in Mubritinib sensitive and resistant primary AML specimens (see cohort in Table S6). Statistical assessment in **a.** and **f.-g.** and **k.** was performed using the Mann-Whitney test. In **l,** the number of Annexin V and Propidium Iodide (PI) double positive cells were considered for statistical assessment and the two-tailed unpaired *t*-test was used. Data in **a.** are represented as mean values with SD, in **b.** and **d.-k.** and **j.** as mean values with SEM and in **k**. as median values.

**Fig. S4, related to Fig. 4: Identification of Mubritinib's target.**

**a.** Schematic overview of the CRISPR whole genome screen carried out in Nalm6-Cas9 clonal cells. **b.** and **c.** Mubritinib dose response assay of OCI-AML3 cells infected with different shRNAs, as indicated. **d**, Inhibitory patterns of Mubritinib (500nM, 6 days) and Oligomycin (10nM, 6 days, ETC complex I inhibitor) in primary AML specimens (see cohort details in Table S7). **e-k**. report the results of cell free enzymatic activity assays. **e,** ETC complex IV activity inhibited in a dose dependent fashion by KCN but **f,** not by Mubritinib. **g.** ETC complex V activity inhibited in a dose dependent fashion by Oligomycin but **h.** not by Mubritinib. **i**, Mubritinib's lack of effect on the activities of ETC complexes II and III compared to Antimycin A used as a positive control. **j.** Rotenone's dose-dependent inhibitory effect on ETC complex I. **k,** Mubritinib's lack of inhibitory effect on ETC complex I activity in a ubiquinone-independent assay using OCI-AML3 cells. **l,**

Effect of Mubritinib treatment (500nM, 20h) on NAD/NADH concentration ratio in OCI-AML3 cells. Data in **b-c. and e-k.** are represented as mean values with SEM and in **l.**, as median values. Statistical assessments in **b.-c** and **l**. were performed using the unpaired two-tailed *t*-test. In **d**, statistical assessments were performed using the Pearson correlation test. In **i**, statistical assessment was performed using the one-way ANOVA test and in **k.,** using the Mann-Whitney test.

**Fig. S5, related to Fig. 5: Characterization of Mubritinib's anti-leukemic activity *in vivo.***
**a.** Flow cytometric analysis of bone marrow cells isolated from the deceased mouse at day 20. Frequency of tomato-positive cells detected **b.** in the bone marrow and **c.** in the spleens of Mubritinib- *versus* vehicle-treated animals at day 29. **d.** Number of tomato-negative nucleated cells detected in the bone marrow (2 hips, 2 femurs and 2 tibias) of Mubritinib- *versus* vehicle-treated animals at day 29. **e.** treatment scheme for the evaluation of Mubritinib's effect on normal hematopoiesis in Pep3B mice. **f.** to **n.** results (in absolute numbers) of FACS analysis of bone marrow cells isolated from 2 hips, 2 femurs and 2 tibias of Mubritinib- *versus* vehicle-treated Pep3B mice at day 29, as indicated. **o.** to **t.,** results of the analysis of blood samples isolated from Mubritinib- *versus* vehicle-treated Pep3B mice at day 29, as indicated. Statistical assessment in **b.-d.** and **f.** to **n.** was performed using the Mann-Whitney test and in **o.-t.** using the unpaired *t*-test. Data in **b.-d.** and **f.-o.** are represented as median values.

**Table S1, related to Fig. 1**
**Primary screen cohort details.**

**Table S2, related to Fig. 1**
**Secondary screen cohort details.**

**Table S3, related to Fig. 2**
**List of clinical and mutational parameters and associated values used in Fig. 2a.**

**Table S4, related to Fig. 2**
**List of most differentially expressed genes in Mubritinib-sensitive (EC50<374nM, n=100) *versus* –resistant (EC50≥374nM, n=100) specimens.** Criteria used: log(fold change)>0.8 (*i.e.* fold change in average RPKM >6) and RPKM>0.1. *HOX*-network genes are highlighted in bold.

**Table S5, related to Fig. 3**
**List of proteins detected by LC/MS-MS in OCI-AML3 cells.**

**Table S6, related to Fig. 3 and S3**
**Cohort details related to Fig. 3s-t and S3j-k.**

**Table S7, related to Fig. 4 and S4**
**Cohort details related to Fig. 4g and S4c.**

**Table S8, related to Fig. 4**
**Structure of analogs included in Fig. 4j.**

**Read file 1: Sequencing results of day 0 of chemogenomic screen**

**Read file 2: Sequencing results of Mubritinib-treated cells**

**Read file 3: Sequencing results of DMSO-treated cells**

**Fig. S1, related to Fig. 1:**
**Clinical, mutational and transcriptional landscape of poor outcome AML specimens.**

**a**

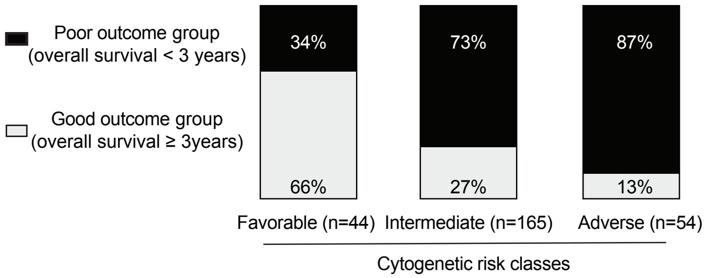


**b**

| | | Poor outcome specimens (n=183) | Good outcome specimens (n=80) | Significant p-values after Bonferroni correction |
|---|---|---|---|---|
| **age** | | **58 (20-78)** | 45 (17-71) | **<0.0001** |
| WBC ($x10^9$/L) | | 42 (1.3-361) | 37 (1-447) | n.s. |
| Gender | | | | |
| Male | | 96 (53%) | 45 (56%) | n.s. |
| Female | | 87(47%) | 35 (44%) | n.s. |
| **Relapse** | | **93 (51%),** NA=59(32%) | 21 (26%), NA=0 | **$p=5x10^{-12}$** |
| Cytogenetic risk | | | | |
| **Favorable** | | 15 (8%) | **29 (36%)** | **<0.0001** |
| Intermediate | | 121 (66%) | 44 (55%) | n.s. |
| **Adverse** | | **47 (26%)** | 7 (9%) | **p=0.0009** |
| FAB | | | | |
| M0 | | 9 (5%) | 2 (3%) | n.s. |
| M1 | | 64 (35%) | 19 (24%) | n.s. |
| M2 | | 22 (12%) | 18 (23%) | n.s. |
| M3 | | 0 (0%) | 0 (0%) | n.s. |
| M4 | | 19 (10%) | 5 (6%) | n.s. |
| M4Eo | | 8 (4%) | 13 (16%) | n.s. |
| M5 | | 3 (2%) | 2 (3%) | n.s. |
| M5A | | 18 (10%) | 5 (6%) | n.s. |
| M5B | | 11 (6%) | 2 (3%) | n.s. |
| M6 | | 1 (1%) | 2 (3%) | n.s. |
| M6A | | 1 (1%) | 0 (0%) | n.s. |
| M6B | | 1 (1%) | 0 (0%) | n.s. |
| M7 | | 0 (0%) | 1 (1%) | n.s. |
| NC | | 26 (14%) | 11 (14%) | n.s. |
| Genetic group | | | | |
| **CBF** | **t(8;21)** | 4 (2%) | **12 (15%)** | **0.0002** / **$4x10^{-8}$** |
| | **inv(16)** | 11 (6%) | **17 (21%)** | **0.0004** |
| Normal karyotype | | 76 (42%) | 28 (35%) | n.s. |
| Intermediate abn. K | | 28 (15%) | 10 (13%) | n.s. |
| NUP98-NSD1 | | 3 (2%) | 2 (3%) | n.s. |
| Trisomy/tetrasomy 8 | | 6 (3%) | 2 (3%) | n.s. |
| MLL | | 18 (10%) | 6 (8%) | n.s. |
| Monosomy 5 | | 4 (2%) | 0 (0%) | n.s. |
| Complex | | 27 (15%) | 2 (3%) | n.s. |
| EVI1 | | 4 (2%) | 0 (0%) | n.s. |
| Other | | 2 (1%) | 1 (1%) | n.s. |

**c**

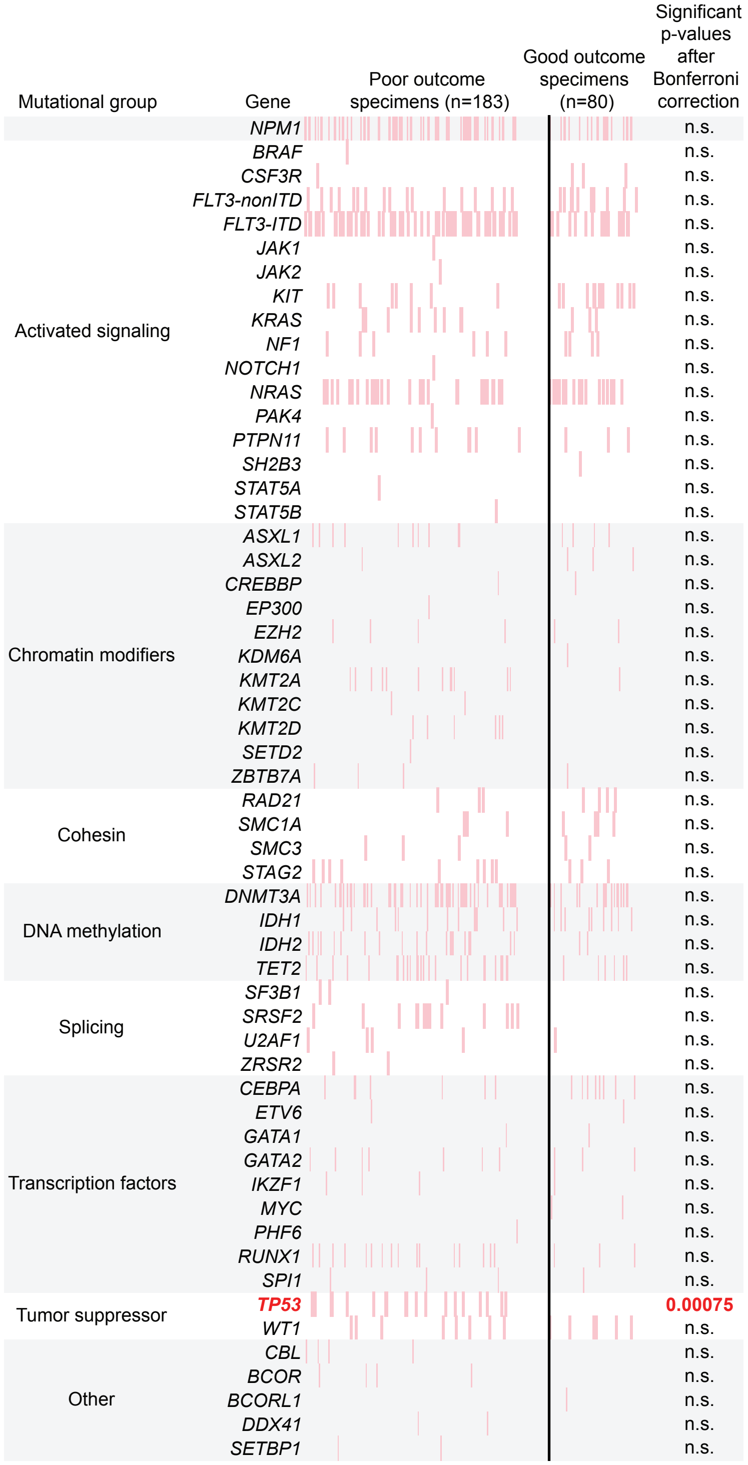


**d**

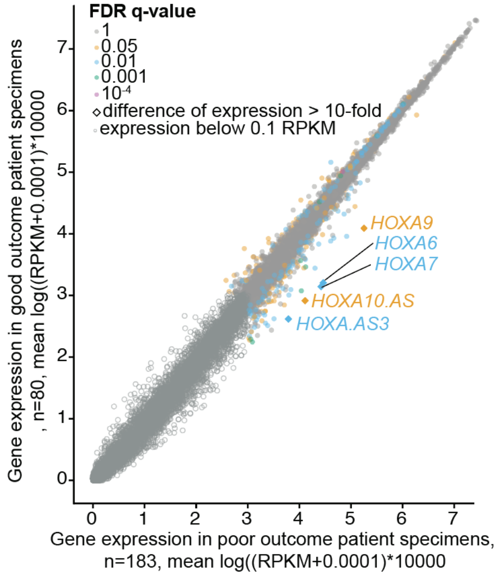

**Fig. S2, related to Fig. 2:**
**Clinical and mutational landscape of Mubritinib sensitive AML specimens.**

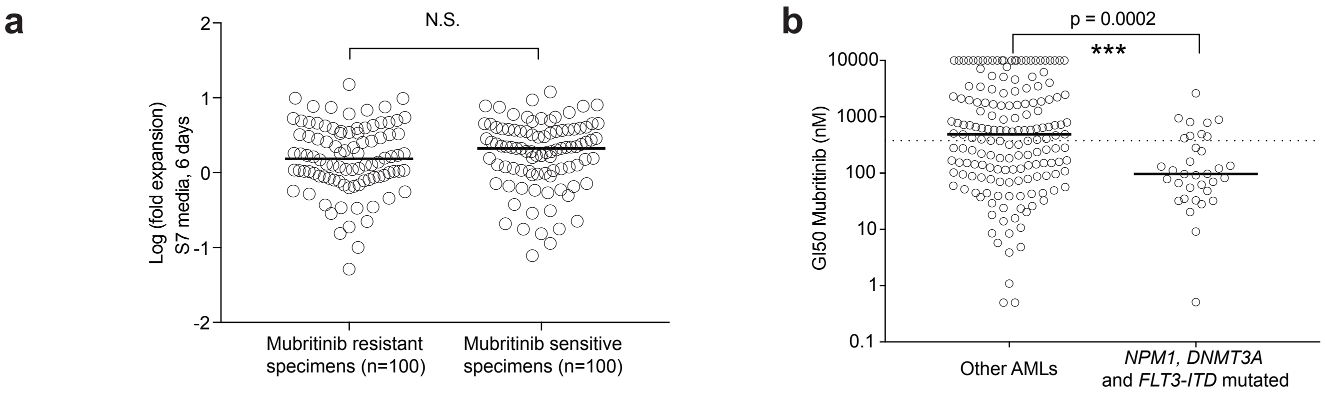

**Fig. S3, related to Fig. 3:**
**Dissection of Mubritinib's mechanism of action.**

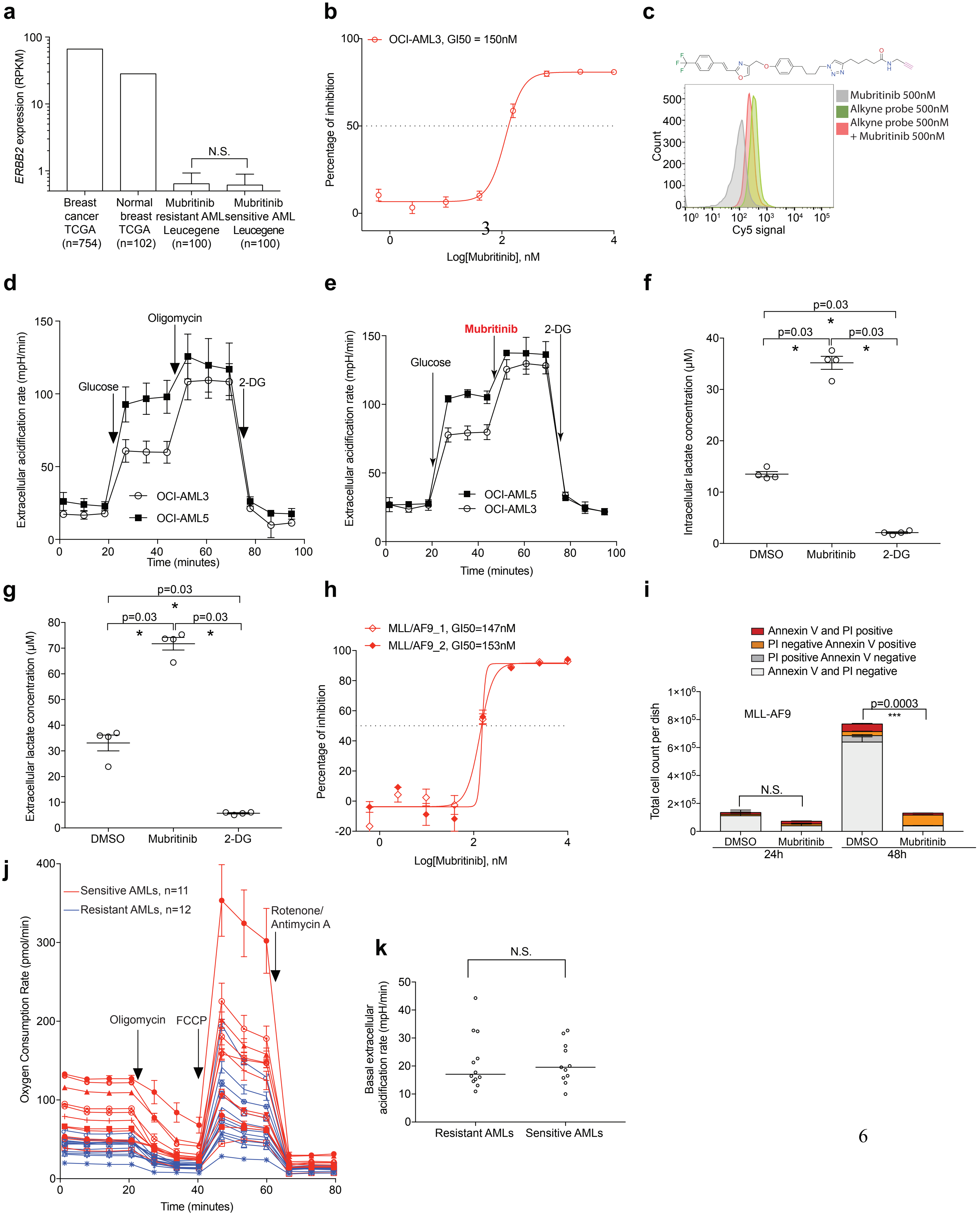

**Fig. S4, related to Fig. 4:**
**Identification of Mubritinib's target.**

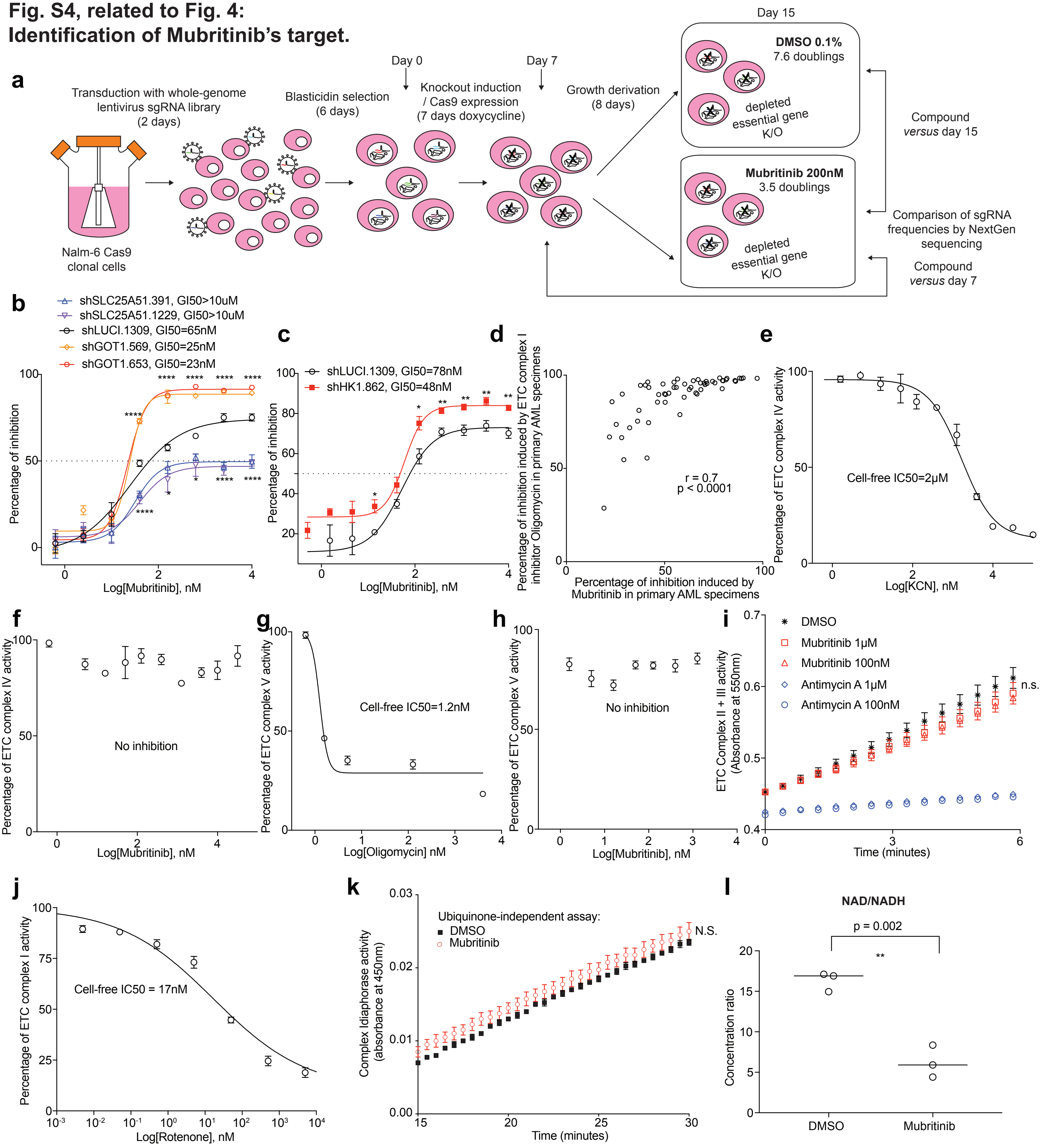

**Fig. S5, related to Fig. 5**
**Characterization of Mubritinib's anti-leukemic activity *in vivo*.**

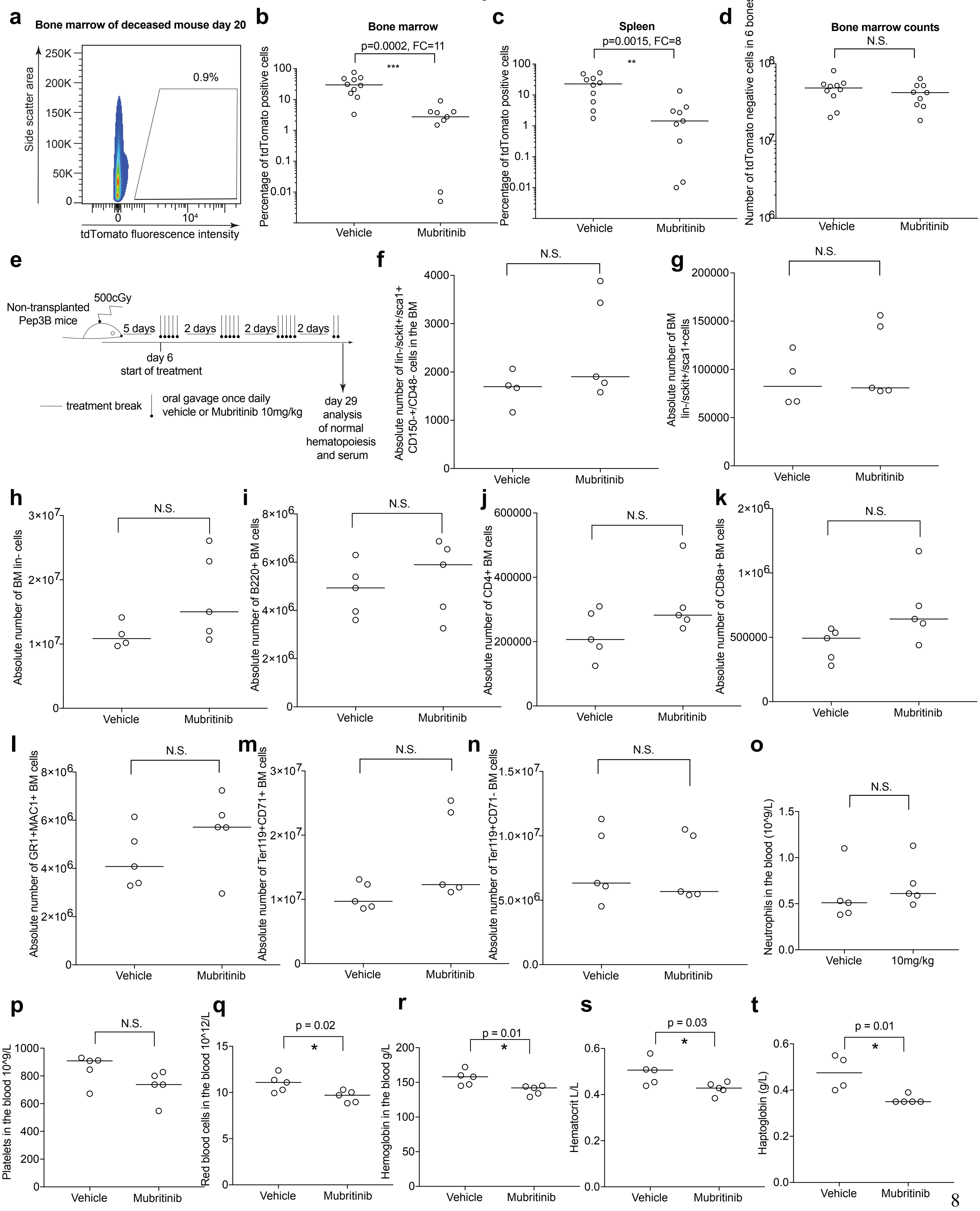

### Table S1, related to Fig. 1
### Primary screen cohort details.

| Patient ID | Sex | Age | Cytogenetic risk class | FAB | WBC (x109/ L) | Genetic subgroup | GI50 Mubritinib (nM) | GI50 Lapatinib (nM) |
|---|---|---|---|---|---|---|---|---|
| 02H017 | F | 50 | adverse cytogenetics | AML-M0 | 51.1 | MLL translocations (+MLL FISH positive) (Irrespective of additional cytogenetic abnormalities) | 658 | ≥10000 |
| 03H109 | M | 29 | favorable cytogenetics | AML-M4Eo | 182 | nv(16)(p13.1q22)/t(16;16)(p13.1;q22)/CBFB-MYH11 (Irrespective of additional cytogenetic abnormalities | 528 | ≥10000 |
| 04H030 | M | 50 | favorable cytogenetics | AML-M4Eo | 17 | nv(16)(p13.1q22)/t(16;16)(p13.1;q22)/CBFB-MYH11 (Irrespective of additional cytogenetic abnormalities | ≥10000 | ≥10000 |
| 05H066 | F | 26 | adverse cytogenetics | AML-M4 | 26.6 | MLL translocations (+MLL FISH positive) (Irrespective of additional cytogenetic abnormalities) | 124 | ≥10000 |
| 05H113 | F | 63 | favorable cytogenetics | AML-M4Eo | 20 | nv(16)(p13.1q22)/t(16;16)(p13.1;q22)/CBFB-MYH11 (Irrespective of additional cytogenetic abnormalities | 68 | ≥10000 |
| 05H136 | M | 66 | favorable cytogenetics | AML-M4Eo | 40.5 | nv(16)(p13.1q22)/t(16;16)(p13.1;q22)/CBFB-MYH11 (Irrespective of additional cytogenetic abnormalities | 277 | ≥10000 |
| 06H088 | M | 28 | adverse cytogenetics | AML-M1 | 32.7 | MLL translocations (+MLL FISH positive) (Irrespective of additional cytogenetic abnormalities) | 184 | ≥10000 |
| 06H115 | M | 40 | favorable cytogenetics | AML-M2 | 18 | nv(16)(p13.1q22)/t(16;16)(p13.1;q22)/CBFB-MYH11 (Irrespective of additional cytogenetic abnormalities | 1034 | ≥10000 |
| 06H117 | M | 58 | adverse cytogenetics | AML-M4 | 87.4 | MLL translocations (+MLL FISH positive) (Irrespective of additional cytogenetic abnormalities) | 536 | ≥10000 |
| 06H144 | F | 71 | intermediate cytogenetics | AML-M1 | 56.1 | Normal karyotype | 877 | ≥10000 |
| 07H099 | F | 58 | favorable cytogenetics | Not classifiable by FAB criteria | 61.5 | nv(16)(p13.1q22)/t(16;16)(p13.1;q22)/CBFB-MYH11 (Irrespective of additional cytogenetic abnormalities | 3665 | ≥10000 |
| 07H135 | M | 67 | intermediate cytogenetics | AML-M1 | 106.7 | Normal karyotype | 254 | 5791 |
| 07H160 | F | 67 | adverse cytogenetics | AML-M1 | 92.2 | MLL translocations (+MLL FISH positive) (Irrespective of additional cytogenetic abnormalities) | 164 | ≥10000 |
| 08H021 | M | 60 | intermediate cytogenetics | AML-M5A | 97.2 | MLL translocations (+MLL FISH positive) (Irrespective of additional cytogenetic abnormalities) | 749 | ≥10000 |
| 08H081 | F | 39 | favorable cytogenetics | AML-M4Eo | 53.7 | nv(16)(p13.1q22)/t(16;16)(p13.1;q22)/CBFB-MYH11 (Irrespective of additional cytogenetic abnormalities | ≥10000 | ≥10000 |
| 08H099 | M | 75 | favorable cytogenetics | AML-M4Eo | 104.5 | nv(16)(p13.1q22)/t(16;16)(p13.1;q22)/CBFB-MYH11 (Irrespective of additional cytogenetic abnormalities | ≥10000 | ≥10000 |
| 09H010 | M | 20 | intermediate cytogenetics | AML-M5A | 1.8 | MLL translocations (+MLL FISH positive) (Irrespective of additional cytogenetic abnormalities) | 140 | ≥10000 |
| 09H016 | F | 64 | favorable cytogenetics | AML-M4Eo | 61.7 | nv(16)(p13.1q22)/t(16;16)(p13.1;q22)/CBFB-MYH11 (Irrespective of additional cytogenetic abnormalities | ≥10000 | ≥10000 |
| 09H018 | M | 56 | adverse cytogenetics | AML-M0 | 101.8 | MLL translocations (+MLL FISH positive) (Irrespective of additional cytogenetic abnormalities) | 6716 | ≥10000 |
| 09H031 | F | 54 | intermediate cytogenetics | AML-M1 | 48.9 | Normal karyotype | 191 | ≥10000 |
| 09H066 | M | 68 | favorable cytogenetics | AML-M4 | 20.7 | nv(16)(p13.1q22)/t(16;16)(p13.1;q22)/CBFB-MYH11 (Irrespective of additional cytogenetic abnormalities | 556 | ≥10000 |
| 09H083 | F | 63 | intermediate cytogenetics | AML-M1 | 65.3 | Normal karyotype | 149 | 4525 |
| 09H111 | F | 59 | intermediate cytogenetics | AML-M5B | 46.8 | Normal karyotype | ≥10000 | ≥10000 |
| 09H113 | M | 56 | intermediate cytogenetics | AML-M1 | 68.7 | Normal karyotype | 1238 | ≥10000 |
| 10H008 | M | 37 | favorable cytogenetics | AML-M4Eo | 75.8 | nv(16)(p13.1q22)/t(16;16)(p13.1;q22)/CBFB-MYH11 (Irrespective of additional cytogenetic abnormalities | 439 | ≥10000 |
| 10H031 | F | 25 | adverse cytogenetics | AML-M5B | 202 | MLL translocations (+MLL FISH positive) (Irrespective of additional cytogenetic abnormalities) | 124 | ≥10000 |
| 10H038 | F | 67 | intermediate cytogenetics | AML-M0 | 147.8 | NUP98-NSD1(caryotype normal) | 386 | ≥10000 |
| 10H056 | F | 57 | intermediate cytogenetics | AML-M1 | 6.3 | Normal karyotype | 70 | 6091 |
| 10H092 | F | 69 | intermediate cytogenetics | AML-M1 | 81.7 | Normal karyotype | 97 | ≥10000 |
| 10H127 | M | 36 | intermediate cytogenetics | AML-M5A | 162.3 | MLL translocations (+MLL FISH positive) (Irrespective of additional cytogenetic abnormalities) | 102 | ≥10000 |
| 11H022 | F | 47 | favorable cytogenetics | AML-M4Eo | 35.8 | nv(16)(p13.1q22)/t(16;16)(p13.1;q22)/CBFB-MYH11 (Irrespective of additional cytogenetic abnormalities | 3668 | ≥10000 |
| 11H072 | F | 52 | intermediate cytogenetics | AML-M2 | 32.2 | Normal karyotype | 46 | ≥10000 |
| 11H104 | M | 56 | favorable cytogenetics | AML-M4Eo | 9.6 | nv(16)(p13.1q22)/t(16;16)(p13.1;q22)/CBFB-MYH11 (Irrespective of additional cytogenetic abnormalities | ≥10000 | ≥10000 |
| 11H129 | M | 62 | intermediate cytogenetics | AML-M1 | 322.5 | Intermediate abnormal karyotype (except isolated trisomy/tetrasomy 8) | 133 | ≥10000 |
| 11H179 | M | 22 | favorable cytogenetics | AML-M4Eo | 16.4 | nv(16)(p13.1q22)/t(16;16)(p13.1;q22)/CBFB-MYH11 (Irrespective of additional cytogenetic abnormalities | 1585 | 9560 |
| 12H042 | M | 58 | favorable cytogenetics | AML-M4Eo | 107.6 | nv(16)(p13.1q22)/t(16;16)(p13.1;q22)/CBFB-MYH11 (Irrespective of additional cytogenetic abnormalities | 7358 | ≥10000 |
| 12H165 | M | 56 | favorable cytogenetics | Not classifiable by FAB criteria | 59.3 | nv(16)(p13.1q22)/t(16;16)(p13.1;q22)/CBFB-MYH11 (Irrespective of additional cytogenetic abnormalities | 686 | ≥10000 |

### Table S3, related to Fig. 2
### List of clinical and mutational parameters and associated values used in Fig. 2a.

| | | Mubritinib sensitive specimens (n=100) | Mubritinib resistant specimens (n=100) | Significant p-values after Bonferroni correction, odds ratio, and [95% confidence interval] | |
|---|---|---|---|---|---|
| | Cytogenetic risk | | | | |
| | **Favorable** | 4 | **33** | **4.4E-08** | **11.7, [4.5;Inf]** |
| | **Intermediate** | **83** | 44 | **6.6E-09** | **6.2, [3.4;Inf]** |
| | Adverse | 13 | 23 | n.s. | |
| | Und | 1 | 0 | n.s. | |
| | FAB | | | | |
| | M0 | 1 | 7 | n.s. | |
| | M1 | 40 | 26 | n.s. | |
| | M2 | 16 | 12 | n.s. | |
| | M3 | 0 | 0 | n.s. | |
| | M4 | 11 | 9 | n.s. | |
| | M4Eo | 3 | 16 | n.s. | |
| | M5 | 2 | 2 | n.s. | |
| | M5A | 7 | 9 | n.s. | |
| | M5B | 7 | 4 | n.s. | |
| | M6 | 0 | 0 | n.s. | |
| | M6A | 1 | 0 | n.s. | |
| | M6B | 0 | 0 | n.s. | |
| | M7 | 0 | 0 | n.s. | |
| | NC | 12 | 15 | n.s. | |
| | Genetic subtype | | | | |
| **CBF** | t(8;21) | 1 | 11 | n.s. | **4.4E-08, 11.7, [4.5;Inf]** |
| **CBF** | **inv(16)** | 3 | **22** | **2.9E-05** | **9.0, [3.0;Inf]** |
| | **Normal karyotype** | **59** | 25 | **8.9E-07** | **4.3, [2.5;Inf]** |
| | Intermediate abn. karyotype | 17 | 11 | n.s. | |
| | NUP98-NSD1 | 2 | 2 | n.s. | |
| | Trisomy/tetrasomy 8 | 2 | 1 | n.s. | |
| | MLL | 7 | 12 | n.s. | |
| | Monosomy 5 | 1 | 1 | n.s. | |
| | Complex | 7 | 12 | n.s. | |
| | EVI1 | 0 | 3 | n.s. | |
| | Other | 1 | 0 | n.s. | |
| | Mutations | | | | |
| | ***NPM1*** | **52** | 24 | **3.7E-05** | **3.4, [2.0;Inf]** |
| RAS/MAPK signaling pathway | *BRAF* | 0 | 1 | n.s. | |
| RAS/MAPK signaling pathway | ***KIT*** | 1 | **18** | **1.8E-05** | **21.5, [3.9;Inf]** |
| RAS/MAPK signaling pathway | *KRAS* | 5 | 5 | n.s. | |
| RAS/MAPK signaling pathway | *NF1* | 1 | 4 | n.s. | |
| RAS/MAPK signaling pathway | *NRAS* | 15 | 24 | n.s. | **1.79E-05, 3.4, [2.0;Inf]** |
| RAS/MAPK signaling pathway | *PTPN11* | 6 | 4 | n.s. | |
| RAS/MAPK signaling pathway | *TP53* | 5 | 7 | n.s. | |
| Activated signaling other than RAS/MAPK signaling pathway | *CSF3R* | 2 | 1 | n.s. | |
| Activated signaling other than RAS/MAPK signaling pathway | *FLT3-nonITD* | 10 | 17 | n.s. | |
| Activated signaling other than RAS/MAPK signaling pathway | ***FLT3-ITD*** | **43** | 19 | **2.0E-04** | **3.2, [1.8;Inf]** |
| Activated signaling other than RAS/MAPK signaling pathway | *JAK1* | 0 | 1 | n.s. | |
| Activated signaling other than RAS/MAPK signaling pathway | *JAK2* | 1 | 0 | n.s. | |
| Activated signaling other than RAS/MAPK signaling pathway | *NOTCH1* | 0 | 1 | n.s. | |
| Activated signaling other than RAS/MAPK signaling pathway | *PAK4* | 0 | 1 | n.s. | |
| Activated signaling other than RAS/MAPK signaling pathway | *SH2B3* | 0 | 1 | n.s. | |
| Activated signaling other than RAS/MAPK signaling pathway | *STAT5B* | 1 | 0 | n.s. | |
| Chromatin modifiers | *ASXL1* | 4 | 7 | n.s. | |
| Chromatin modifiers | *ASXL2* | 0 | 2 | n.s. | |
| Chromatin modifiers | *CREBBP* | 0 | 1 | n.s. | |
| Chromatin modifiers | *EP300* | 1 | 0 | n.s. | |
| Chromatin modifiers | *EZH2* | 1 | 1 | n.s. | |
| Chromatin modifiers | *KDM6A* | 0 | 1 | n.s. | |
| Chromatin modifiers | *KMT2A* | 9 | 2 | n.s. | |
| Chromatin modifiers | *KMT2C* | 1 | 1 | n.s. | |
| Chromatin modifiers | *KMT2D* | 5 | 2 | n.s. | |
| Chromatin modifiers | *WHSC1* | 1 | 0 | n.s. | |
| Chromatin modifiers | *ZBTB7A* | 1 | 2 | n.s. | |
| Cohesin | *RAD21* | 5 | 1 | n.s. | |
| Cohesin | *SMC1A* | 4 | 3 | n.s. | |
| Cohesin | *SMC3* | 2 | 1 | n.s. | |
| Cohesin | *STAG2* | 7 | 2 | n.s. | |
| **DNA methylation** | ***DNMT3A*** | **47** | 23 | **3.0E-04** | **3.0, [1.7;Inf]** |
| **DNA methylation** | *IDH1* | 12 | 3 | n.s. | |
| **DNA methylation** | *IDH2* | 13 | 8 | n.s. | **1.9E-05, 2.3, [1.6;Inf]** |
| **DNA methylation** | *TET2* | 16 | 10 | n.s. | |
| Splicing | *SF3B1* | 3 | 1 | n.s. | |
| Splicing | *SRSF2* | 9 | 2 | n.s. | |
| Splicing | *U2AF1* | 2 | 1 | n.s. | |
| Splicing | *ZRSR2* | 0 | 1 | n.s. | |
| Transcription factors | *CEBPA* | 13 | 1 | n.s. | |
| Transcription factors | *ETV6* | 2 | 1 | n.s. | |
| Transcription factors | *GATA1* | 1 | 0 | n.s. | |
| Transcription factors | *GATA2* | 6 | 3 | n.s. | |
| Transcription factors | *PHF6* | 1 | 0 | n.s. | |
| Transcription factors | *RUNX1* | 9 | 5 | n.s. | |
| Transcription factors | *SPI1* | 0 | 3 | n.s. | |
| Tumor suppressor | *WT1* | 8 | 5 | n.s. | |
| Other | *CBL* | 1 | 3 | n.s. | |
| Other | *BCOR* | 1 | 3 | n.s. | |
| Other | *BCORL1* | 0 | 1 | n.s. | |
| Other | *DDX41* | 2 | 0 | n.s. | |
| Other | *SETBP1* | 0 | 2 | n.s. | |

#### Table S4, related to Fig. 2
#### List of most differentially expressed genes in Mubritinib-sensitive *versus* resistant specimens.

| Gene | FDR q-value | Expression (*) in Mubritinib resistant AML | Expression (*) in Mubritinib sensitive AML |
|---|---|---|---|
| *SNORD116.4* | 0.0002 | 3.9 | 3.0 |
| *SNORD116.24* | 0.0008 | 3.3 | 2.4 |
| *ZNF521* | 0.0031 | 4.5 | 3.6 |
| *MSLN* | 0.0001 | 3.9 | 2.6 |
| *S100A16* | 0.0011 | 3.5 | 2.5 |
| *SNORD116.20* | 0.0003 | 3.4 | 2.4 |
| *KIRREL* | 0.0001 | 3.0 | 2.0 |
| *PRG3* | 0.0015 | 3.9 | 3.0 |
| *SNORD116.21* | 0.0016 | 3.4 | 2.5 |
| ***HOXA5*** | 0.0065 | 4.2 | 5.0 |
| ***HOXB5*** | 0.0074 | 3.1 | 4.0 |
| ***HOXB9*** | 0.0078 | 2.3 | 3.2 |
| ***HOXA11*** | 0.0206 | 2.5 | 3.3 |
| *COL4A5* | 0.0006 | 2.8 | 4.0 |
| *PRDM16* | 0.0024 | 2.7 | 3.5 |
| *BEND6* | 0.0024 | 2.8 | 3.6 |
| *LINC00982* | 0.0025 | 2.1 | 3.1 |
| *NKX2.3* | 0.0002 | 2.4 | 3.9 |
| *ANKRD18B* | 0.0001 | 2.5 | 3.4 |
| *MIR4740* | 0.0011 | 3.4 | 4.3 |
| ***HOXA7*** | 0.0043 | 3.6 | 4.5 |
| ***HOXB.AS3*** | 0.0004 | 2.7 | 4.1 |
| ***HOXA.AS3*** | 0.0051 | 2.9 | 3.9 |
| ***HOXA6*** | 0.0055 | 3.5 | 4.5 |

(*) Expression in Log((RPKM+0.0001)*10000)

**Table S6, related to Fig. 3**
**Cohort details related to Fig. 3s-t and S3j-k.**

| Patient ID | Sensitive (S) or Resistant (R) group | GI0 Mubritinib (nM) | Genetic subgroup | Cytogenetic risk class |
|---|---|---|---|---|
| 04H133 | S | 9 | Normal karyotype | intermediate cytogenetics |
| 05H008 | S | 18 | Normal karyotype | intermediate cytogenetics |
| 06H011 | S | 59 | Normal karyotype | intermediate cytogenetics |
| 07H042 | S | 67 | Normal karyotype | intermediate cytogenetics |
| 10H101 | S | 32 | Normal karyotype | intermediate cytogenetics |
| 10H166 | S | 55 | Normal karyotype | intermediate cytogenetics |
| 11H157 | S | 48 | Normal karyotype | intermediate cytogenetics |
| 11H234 | S | 26 | Normal karyotype | intermediate cytogenetics |
| 12H173 | S | 43 | Intermediate abnormal | intermediate cytogenetics |
| 13H150 | S | 29 | Complex | adverse cytogenetics |
| 14H007 | S | 62 | Normal karyotype | intermediate cytogenetics |
| 03H112 | R | > 10,000 | inv(16) | favorable cytogenetics |
| 05H118 | R | 7723 | t(8;21) | favorable cytogenetics |
| 07H148 | R | > 10,000 | Complex | adverse cytogenetics |
| 08H081 | R | 5313 | inv(16) | favorable cytogenetics |
| 08H099 | R | > 10,000 | inv(16) | favorable cytogenetics |
| 09H016 | R | 7159 | inv(16) | favorable cytogenetics |
| 09H111 | R | 5106 | Normal karyotype | intermediate cytogenetics |
| 10H119 | R | > 10,000 | t(8;21) | favorable cytogenetics |
| 10H174 | R | > 10,000 | Complex | adverse cytogenetics |
| 12H042 | R | > 10,000 | inv(16) | favorable cytogenetics |
| 12H165 | R | > 10,000 | inv(16) | favorable cytogenetics |
| 12H170 | R | > 10,000 | Complex | adverse cytogenetics |

#### Table S7, related to Fig. 4
#### Cohort details related to Fig. 4g and S4c.

| Patient ID | Sex | Age | Cytogenetic risk class | FAB | WBC (x109/ L) | Genetic subgroup |
|---|---|---|---|---|---|---|
| 05H094 | M | 35 | intermediate cytogenetics | AML-M5B | 98.1 | Normal karyotype |
| 05H186 | M | 73 | intermediate cytogenetics | Not classifiable by FAB criteria | 99.2 | Normal karyotype |
| 06H021 | F | 49 | intermediate cytogenetics | AML-M2 | 68.2 | Normal karyotype |
| 06H088 | M | 28 | adverse cytogenetics | AML-M1 | 32.7 | MLL translocations (+MLL FISH positive) (Irrespective of additional cytogenetic abnormalities) |
| 06H089 | M | 58 | intermediate cytogenetics | AML-M1 | 43.5 | Intermediate abnormal karyotype (except isolated trisomy/tetrasomy 8) |
| 06H117 | M | 58 | adverse cytogenetics | AML-M4 | 87.4 | MLL translocations (+MLL FISH positive) (Irrespective of additional cytogenetic abnormalities) |
| 06H143 | M | 73 | intermediate cytogenetics | AML-M4 | 153 | Intermediate abnormal karyotype (except isolated trisomy/tetrasomy 8) |
| 06H146 | M | 45 | intermediate cytogenetics | AML-M2 | 61.9 | Normal karyotype |
| 07H003 | F | 25 | adverse cytogenetics | AML-M4 | 46 | MLL translocations (+MLL FISH positive) (Irrespective of additional cytogenetic abnormalities) |
| 07H069 | M | 55 | intermediate cytogenetics | Not classifiable by FAB criteria | 80 | Intermediate abnormal karyotype (except isolated trisomy/tetrasomy 8) |
| 07H089 | F | 37 | intermediate cytogenetics | AML-M5 | 76.5 | Normal karyotype |
| 07H125 | F | 66 | intermediate cytogenetics | AML-M1 | 45.5 | Normal karyotype |
| 07H135 | M | 67 | intermediate cytogenetics | AML-M1 | 106.7 | Normal karyotype |
| 08H018 | M | 65 | adverse cytogenetics | Not classifiable by FAB criteria | 19.9 | Complex (3 and more chromosomal abnormalities) |
| 08H021 | M | 60 | intermediate cytogenetics | AML-M5A | 97.2 | MLL translocations (+MLL FISH positive) (Irrespective of additional cytogenetic abnormalities) |
| 08H049 | M | 69 | intermediate cytogenetics | AML-M5B | 69 | NUP98-NSD1(normal karyotype) |
| 08H082 | F | 43 | intermediate cytogenetics | AML-M1 | 102.7 | Normal karyotype |
| 08H085 | F | 49 | adverse cytogenetics | AML-M1 | 268 | MLL translocations (+MLL FISH positive) (Irrespective of additional cytogenetic abnormalities) |
| 08H112 | M | 52 | intermediate cytogenetics | Not classifiable by FAB criteria | 28.3 | Normal karyotype |
| 08H113 | M | 53 | intermediate cytogenetics | AML-M1 | 36.6 | Normal karyotype |
| 09H016 | F | 64 | favorable cytogenetics | AML-M4Eo | 61.7 | inv(16)(p13.1q22)/t(16;16)(p13.1;q22)/CBFB-MYH11 (Irrespective of additional cytogenetic abnormalities) |
| 09H026 | F | 73 | intermediate cytogenetics | AML-M1 | 96.8 | Normal karyotype |
| 09H031 | F | 54 | intermediate cytogenetics | AML-M1 | 48.9 | Normal karyotype |
| 09H046 | M | 37 | adverse cytogenetics | Not classifiable by FAB criteria | 52 | Monosomy17/del17p (less than 3 chromosomal abnormalities) |
| 09H054 | F | 54 | adverse cytogenetics | Not classifiable by FAB criteria | 80.3 | Complex (3 and more chromosomal abnormalities) |
| 09H090 | F | 44 | intermediate cytogenetics | AML-M1 | 73.4 | Intermediate abnormal karyotype (except isolated trisomy/tetrasomy 8) |
| 09H106 | M | 74 | intermediate cytogenetics | AML-M5B | 90 | Normal karyotype |
| 09H115 | M | 46 | intermediate cytogenetics | AML-M1 | 101 | Normal karyotype |
| 10H007 | F | 63 | intermediate cytogenetics | AML-M4 | 32.5 | Hyperdiploid numerical abnormalities only |
| 10H008 | M | 37 | favorable cytogenetics | AML-M4Eo | 75.8 | inv(16)(p13.1q22)/t(16;16)(p13.1;q22)/CBFB-MYH11 (Irrespective of additional cytogenetic abnormalities) |
| 10H029 | M | 81 | intermediate cytogenetics | AML-M5B | 87.6 | Intermediate abnormal karyotype (except isolated trisomy/tetrasomy 8) |
| 10H063 | F | 41 | adverse cytogenetics | AML-M4 | 138.1 | EVI1 rearrangements (+EVI1 FISH positive) (Irrespective of additional cytogenetic abnormalities) |
| 10H109 | F | 65 | intermediate cytogenetics | AML-M1 | 318 | Intermediate abnormal karyotype (except isolated trisomy/tetrasomy 8) |
| 10H113 | M | 77 | adverse cytogenetics | AML-M1 | 71.3 | Complex (3 and more chromosomal abnormalities) |
| 10H115 | M | 43 | intermediate cytogenetics | AML-M1 | 21.6 | Normal karyotype |
| 10H127 | M | 36 | intermediate cytogenetics | AML-M5A | 162.3 | MLL translocations (+MLL FISH positive) (Irrespective of additional cytogenetic abnormalities) |
| 10H161 | F | 79 | intermediate cytogenetics | AML-M5B | 248 | Trisomy/tetrasomy 8 (isolated) |
| 10H174 | F | 59 | adverse cytogenetics | AML-M4 | 112.7 | Complex (3 and more chromosomal abnormalities) |
| 11H006 | F | 26 | intermediate cytogenetics | AML-M5A | 33 | Normal karyotype |
| 11H008 | M | 76 | intermediate cytogenetics | Not classifiable by FAB criteria | 86.2 | Intermediate abnormal karyotype (except isolated trisomy/tetrasomy 8) |
| 11H019 | F | 78 | intermediate cytogenetics | AML-M1 | 61.6 | Intermediate abnormal karyotype (except isolated trisomy/tetrasomy 8) |
| 11H046 | M | 77 | intermediate cytogenetics | AML-M0 | 48.8 | Normal karyotype |
| 11H138 | M | 71 | intermediate cytogenetics | AML-M0 | 24 | Hyperdiploid numerical abnormalities only |
| 11H157 | M | 62 | intermediate cytogenetics | AML-M4 | 56.7 | Normal karyotype |
| 11H160 | F | 36 | intermediate cytogenetics | AML-M4 | 207.4 | NUP98-NSD1(normal karyotype) |
| 12H030 | M | 78 | intermediate cytogenetics | AML-M0 | 159.5 | Intermediate abnormal karyotype (except isolated trisomy/tetrasomy 8) |
| 12H106 | M | 67 | adverse cytogenetics | Not classifiable by FAB criteria | 166.8 | Complex (3 and more chromosomal abnormalities) |
| 12H171 | F | 67 | intermediate cytogenetics | AML-M1 | 86.7 | Normal karyotype |
| 12H173 | F | 66 | intermediate cytogenetics | AML-M1 | 206.1 | Intermediate abnormal karyotype (except isolated trisomy/tetrasomy 8) |
| 13H048 | M | 69 | intermediate cytogenetics | AML-M0 | 48.9 | Normal karyotype |
| 13H073 | F | 87 | intermediate cytogenetics | AML-M1 | 357 | Normal karyotype |
| 13H114 | M | 58 | intermediate cytogenetics | Not classifiable by FAB criteria | 68.4 | Normal karyotype |
| 13H141 | M | 83 | adverse cytogenetics | AML-M5A | 55.8 | Complex (3 and more chromosomal abnormalities) |
| 13H150 | M | 60 | adverse cytogenetics | AML-M1 | 62 | Complex (3 and more chromosomal abnormalities) |
| 14H001 | F | 40 | intermediate cytogenetics | AML-M1 | 89.2 | Normal karyotype |
| 14H019 | M | 42 | intermediate cytogenetics | AML-M0 | 19.5 | Normal karyotype |

### Table S8, related to Fig. 4
### Structure of analogs included in Fig. 4j.

| Analog number | Structure, red highlight indicates conserved part compared to Mubritinib | ETC complex I cell free IC50 (nM) | GI50 (nM) in OCI-AML3 | GI50 (nM) in MLL-AF9 |
|---|---|---|---|---|
| 1 | 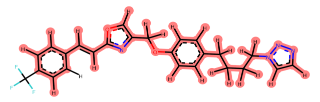 | 26996 | ≥10000 | 7597 |
| 2 | 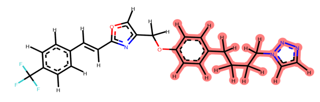 | ≥50000 | ≥10000 | ≥10000 |
| 3 | 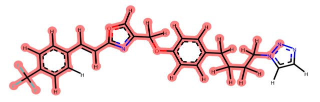 | 19961 | ≥10000 | ≥10000 |
| 4 | 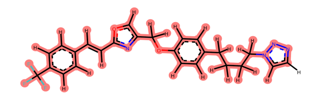 | 7717 | 5476 | 7023 |
| 5 | 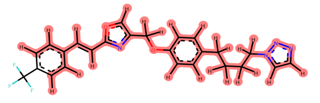 | 1931 | 531 | 580 |
| 6 | 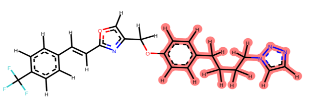 | 1200 | 1084 | 382 |
| 7 | 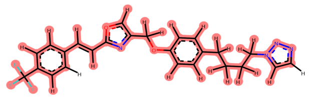 | 703 | 124 | 20 |
| 8 | 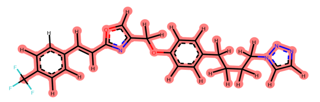 | 518 | 37 | 65 |
| 9 | 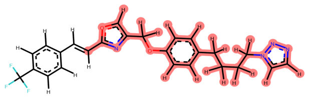 | 140 | 31 | 11 |
| 10 | 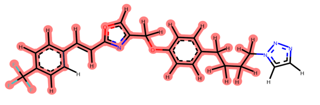 | 114 | 18 | 51 |
| 11 | 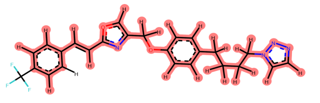 | 81 | 44 | 66 |
| 12 | 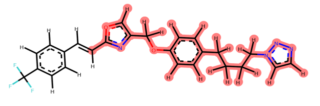 | 52 | 35 | 6 |
| 13 | 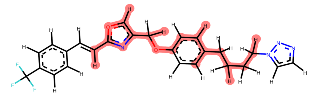 | 45 | 22 | 9 |
| 14 | 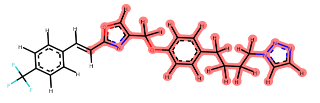 | 39 | 21 | 9 |
| 15 | 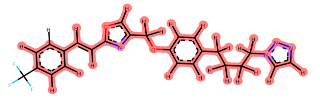 | 30 | 52 | 23 |
